# The effectiveness of pseudomagic traits in promoting premating isolation

**DOI:** 10.1101/2023.01.19.524751

**Authors:** Thomas G. Aubier, Reinhard Bürger, Maria R. Servedio

**Affiliations:** Department of Biology, University of North Carolina at Chapel Hill, Chapel Hill, NC 27599, USA; Department of Mathematics, University of Vienna, 1090 Vienna, Austria

**Keywords:** speciation, mate choice, recombination, secondary contact, third order linkage disequilibrium, mathematical model

## Abstract

Upon the secondary contact of populations, speciation with gene flow is greatly facilitated when the same pleiotropic loci are both subject to divergent ecological selection and induce non-random mating, leading to loci with this fortuitous combination of functions being referred to as “magic trait” loci. We use a population genetics model to examine whether “pseudomagic trait” complexes, composed of physically linked loci fulfilling these two functions, are as efficient in promoting premating isolation as magic traits. We specifically measure the evolution of choosiness, which controls the strength of assortative mating. We show that, surprisingly, pseudomagic trait complexes, and to a lesser extent also physically unlinked loci, can lead to the evolution of considerably stronger assortative mating preferences than do magic traits, provided polymorphism at the involved loci is maintained. This is because assortative mating preferences are generally favored when there is a risk of producing maladapted recombinants, as occurs with nonmagic trait complexes but not with magic traits (since pleiotropy precludes recombination). Contrary to current belief, magic traits may not be the most effective genetic architecture for promoting strong premating isolation. Distinguishing between magic traits and pseudomagic trait complexes is therefore important when inferring their role in premating isolation. This calls for further fine-scale genomic research on speciation genes.

## INTRODUCTION

Assortative mating – the tendency of individuals of similar phenotype to mate more often than expected by chance – has been reported in animals [1, 2] and plants [3, 4], and plays a key role in generating premating reproductive isolation during ecological speciation [5]. When assortative mating is driven by mate choice, the mating rule and nature of the loci underlying assortative mating have important implications for the speciation process [6]. For instance, speciation is less likely to occur if assortative mating is associated with a “preference/trait rule” (i.e., preferences for specific traits used as mating signals) compared to a “matching rule” (i.e., preference for matching mates), partly because speciation under a preference/trait rule requires linkage disequilibrium (i.e., statistical associations between alleles) between more loci [7]. This well-established theoretical result demonstrates that linkage disequilibrium between the loci underlying premating isolation can have a significant impact on speciation with gene flow.

In concert with assortative mating, divergent selection is a key component of speciation with gene flow, as divergent selection drives the incipient species apart [5]. Notably, many models of speciation have shown that across generations, selection generates linkage disequilibrium between ecological loci under divergent selection and loci coding for mating signals [7–10]. All loci subject to divergent selection thus have the potential to strengthen the premating isolation caused by mate choice.

Given that recombination consistently degrades allelic associations, tight physical linkage between the loci underlying premating isolation facilitates the maintenance of strong linkage disequilibrium [7]. In particular, a strong link between divergent ecological selection and assortative mating is guaranteed when the mating signal is itself under divergent ecological selection. Such pleiotropic traits that are both subject to divergent ecological selection and used as the basis of assortative mating (so-called “magic traits” [11]) can facilitate speciation for the simple reason that there is no recombination degrading the link between divergent selection and assortative mating. In other words, pleiotropy of the underlying genes is much more effective than linkage disequilibrium in transmitting the force of divergent ecological selection to the genes causing premating isolation, thereby favoring divergence between incipient species [8, 11, 12]

While many theoretical models of speciation assume magic traits [8, 13–17], it can be very difficult to demonstrate that candidate traits are pleiotropic [6, 18]. In several strong candidate examples of magic traits, it is not yet possible to distinguish between pleiotropy and tight linkage. For instance, the co-localization of quantitative trait loci for traits under divergent selection and traits involved in assortative mating found in *Acyrthosiphon pisum* pea aphids [19] or in *Mimulus* [20] is actually consistent with both possibilities.

Several authors have speculated that “nonmagic” trait complexes, i.e., loci that are coding separately for ecological traits and mating signals, could conceivably mimic the role of magic traits in the speciation process when these loci are tightly physically linked [18, 21, 22]. Strong linkage disequilibrium between a pair of loci – one ecological locus subject to divergent selection and one mating signal locus affecting reproductive isolation – could substitute for the pleiotropic characteristic of true magic traits, assuming that this linkage disequilibrium is strong from the outset of its involvement in speciation (e.g., upon the secondary contact of divergent populations) and assuming that the physical linkage between loci is tight enough [18]. Nonetheless, it is only recently that this intuitive idea has been investigated formally with a population genetics model, with one ecological locus under divergent selection and one locus that acts as a mating signal [23]. Upon secondary contact, such nonmagic trait complexes are actually very effective in promoting divergence, and surprisingly this is also the case if the loci are loosely physically linked, or even if they are physically unlinked (i.e., located on different chromosomes).

Given that nonmagic trait complexes mimic magic traits in terms of the ecological divergence they allow, complexes of physically linked loci coding for separate ecological traits and mating signals have been called “pseudomagic trait” complexes [23]. Although the adjective “pseudomagic” was originally coined to describe gene complexes where tight linkage was indistinguishable from pleiotropy [23], we use it here to describe any physically linked gene complex. Therefore, “nonmagic” trait complexes include pseudomagic trait complexes and freely recombining ecological and mating signal loci. Nevertheless, the question of whether these pseudomagic trait complexes promote speciation in the same way as magic traits cannot be answered on the basis of their effect on divergence alone. The establishment of premating isolation additionally relies on the strength of mate choice (hereafter called ‘choosiness’), which can also be genetically encoded and can therefore be subject to evolution [6]. Linkage disequilibrium plays an important role in the evolution of choosiness (or any ‘one-allele mechanism’ [7]). Yet, magic traits and pseudomagic trait complexes vary notably in terms of the linkage disequilibrium that can be built.

In the case of a magic trait, the evolution of choosiness relies on the linkage disequilibrium between the choosiness locus and the locus encoding the magic trait [8, 13–15]. In contrast, in the case of a pseudomagic trait complex, there are three types of genetic associations which can affect the evolution of choosiness: linkage disequilibria of the choosiness locus with the ecological locus, with the mating signal locus, and with the combination of these two loci (so-called ‘three-way linkage disequilibrium’, or ‘third-order linkage disequilibrium’). The fact that the sources of indirect selection are so different makes it unlikely that the same level of choosiness will evolve with a magic trait and with a pseudomagic trait complex. The evolution of choosiness is thus expected to depend on the genetic architecture underlying ecological and mating signal traits, i.e., on whether choosiness evolves alongside a magic trait or a pseudomagic trait complex.

Using a two-island, three-locus population genetics model, we explore the degree to which pleiotropy (a magic trait) versus varying degrees of physical linkage between an ecological locus and a mating signal locus (pseudomagic trait complexes, and nonmagic traits complexes with physically unlinked loci) can affect the evolution of choosiness upon secondary contact. Our analyses highlight that, although magic traits favor trait divergence and the maintenance of polymorphism, pseudomagic trait complexes, and to a lesser extent physically unlinked loci, can promote stronger choosiness (and premating isolation) than do magic traits. This is because assortative mating preferences are favored when there is a risk of producing maladapted recombinants, as occurs with nonmagic trait complexes but not with magic traits.

## METHODS

### The model

We use a population genetics model to consider a secondary contact scenario in which assortative mating evolves based on three haploid diallelic loci. We implement an ecological locus E, subject to divergent viability selection in both sexes, and a mating signal locus T also expressed in both sexes, on which females base their mate choice using a matching rule (where females prefer to mate with males that have the same mating signal allele as their own; see [6] for a review of the theoretical and empirical literature on phenotype matching). In addition, we implement a choosiness locus C expressed in females. Importantly, choosiness is ecologically neutral and not subject to direct selection. Nevertheless, it undergoes indirect selection via linkage disequilibrium with the other loci (or via linkage disequilibrium with the pleiotropic locus when we consider a magic trait). Importantly, analyses of variants of the model presented here show that our qualitative results are generalizable to diploids and to the evolution of costly choosiness (see Supplementary Information Text).

We assume that two allopatric populations have diverged and that secondary contact occurs between these populations. Each generation first undergoes symmetric migration with rate *m*. Next, divergent selection occurs. The ecological trait encoded by locus E is locally adapted, such that each ecological allele, *E*_1_ and *E*_2_, is favored by viability selection (with selection coefficient *s*) in the population in which it was common in allopatry. Mating follows, during which, females may express different propensities to mate with males. Choosy females prefer to mate with males that match their own mating signal trait encoded by locus T. For instance, a *T*_1_ female is more likely to mate with a *T*_1_ male than a *T*_2_ male upon encounter. Such mate choice generates positive frequency-dependent sexual selection acting on the T locus, because males carrying the locally rare allele at the T locus are unpopular and thus have low mating success [14]. Locus C determines the strength of female preference (hereafter, choosiness), so that *C*_1_ (resp. *C*_2_) females are 1 + *α*_1_ (resp. 1 + *α*_2_) times more likely to mate with a preferred male than with a non-preferred male, if they were to encounter one of each. The expected genotype frequencies in the next generation depend on the probabilities of mating between different genotypes, with the new generation being formed by assuming Mendelian inheritance at all loci. In particular, we assume that gene order is ETC (we relax this assumption in supplementary analyses), and that recombination occurs at a rate *r*_ET_ between loci E and T, and *r*_TC_ between loci T and C. We consider that there is no crossover interference, so that the recombination rate between the E and C loci is determined by the other recombination rates. We assume that loci E and T are initially at maximum linkage disequilibrium in each subpopulation, so that genotypes *E*_1_*T*_2_ and *E*_2_*T*_1_ are initially absent. We can thus model a magic trait by setting *r*_ET_ = 0, a pseudomagic trait complex by assuming 0 *< r*_ET_ *<* 0.5, and physically unlinked E and T loci by considering *r*_ET_ = 0.5.

All recursion equations are detailed in the Supplementary Information Text.

### Migration-selection equilibrium and the maintenance of polymorphism

We initiate the populations assuming that the choosiness allele *C*_1_ is fixed in the two populations; this corresponds to the two-island, two-locus pseudomagic model analyzed in a previous study [23]. Following this previous study, we assume that the two allopatric populations have diverged such that all individuals in population 1 have the *E*_1_ and *T*_1_ alleles and that most individuals in population 2 have the *E*_2_ and *T*_2_ alleles (at a frequency equal to 0.99; other individuals in population 2 have the *E*_1_ and *T*_1_ alleles), thereby avoiding possible artifacts of starting with symmetric initial conditions.

To assess the migration-selection equilibrium, we run numerical iterations until the change in each genotype frequency is less than 10^−9^ per generation. We can thus numerically assess the allelic frequencies and the linkage disequilibria between loci (i.e., statistical associations of alleles at different loci) at migration-selection equilibrium. As a check, the assumption that there is no variation in choosiness allows us to reproduce results of the previous study mentioned earlier [23], including the finding, important for the current study, that divergence at the ecological and mating signal loci, E and T, peaks at intermediate choosiness values (Fig. 1). Indeed, very high choosiness causes rare, very choosy females to mate with rare males in proportion to their frequency, resulting in the loss of positive frequency-dependent sexual selection and thus the reduction or loss of mating signal divergence [16]. Additionally, polymorphism at the T locus can be maintained, in particular for a low recombination rate between the E and T loci (low *r*_ET_) and for an intermediate choosiness value (intermediate *α*_1_; remember that we assume here that allele *C*_2_ is absent) (Fig. 1; as in [23]). Importantly, under random mating (*α*_1_ = 0), we observe that polymorphism at the T locus can be maintained only for *r*_ET_ = 0; with random mating and *r*_ET_ *>* 0, the T locus is neutral and polymorphism can thus not be maintained at it [23].

**Figure 1:**
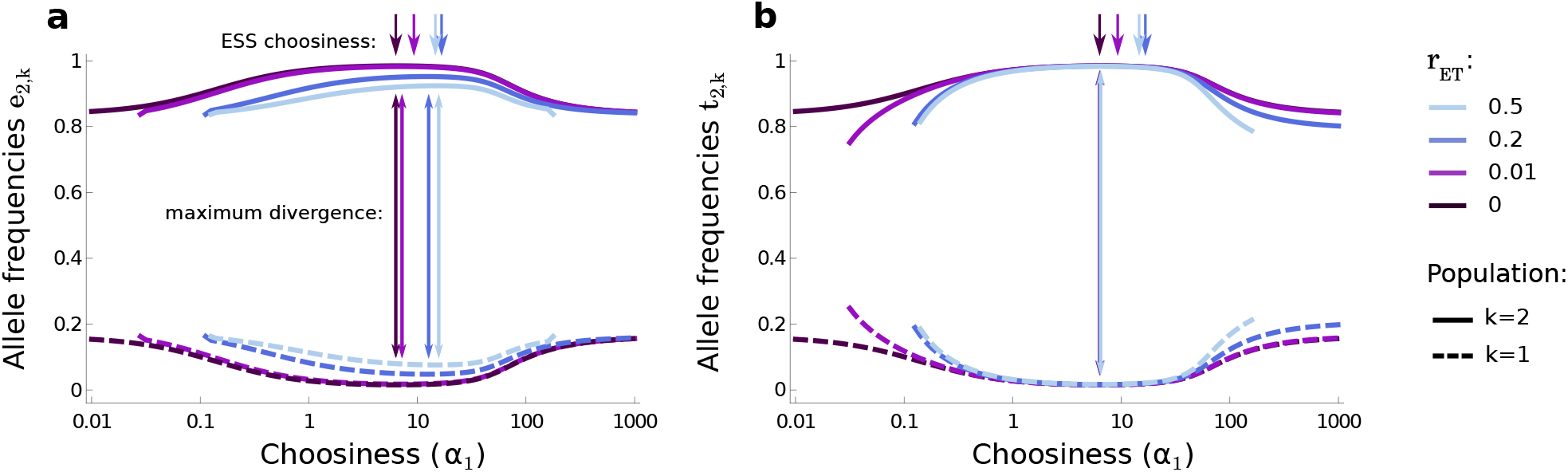
Population divergence depending on genetic architecture and choosiness. We represent the equilibria reached in the two populations after secondary contact for a range of values of choosiness *α*_1_ (assuming that allele *C*_2_ is absent) and recombination rates *r*_ET_, with initial maximum linkage disequilibrium between the E and T loci. We do not represent cases where polymorphism is lost at the T locus (for very low or very high choosiness, as shown where lines end, where frequencies actually fall to zero). Panel **a** shows the equilibrium frequencies *e*_2,*k*_ of allele *E*_2_ in each population *k*. Panel **b** shows the equilibrium frequencies *t*_2,*k*_ of allele *T*_2_ in each population *k*. Double-headed arrows are placed at the choosiness value maximizing divergence at the E and T loci in panel **a** and **b**, respectively. Single-headed arrows correspond to ESS choosiness values that are favored for *r*_TC_ = 0.01 (see Fig. 2). In panel **b**, the double-headed arrows overlap. Note that the divergence-maximizing choosiness (double-headed arrows) and the ESS choosiness (single-headed arrows) do not have the same value, unless *r*_ET_ = 0; this means that additional evolutionary forces, specific to the case where *r*_ET_ *>* 0 and which are the focus of our study, come into play. See ref. [23] for explanations on the divergence pattern. Here, *m* = 0.01, *s* = 0.05.

We next determine the lowest choosiness value, *α*_poly_, that allows the maintenance of polymorphism at both the E and T loci. Using numerical methods detailed in the *Mathematica* notebook, we determine *α*_poly_ with a precision of 10^−3^. Notably, if *α*_poly_ *>* 0, higher choosiness cannot evolve from random mating (*α*_1_ = 0), assuming that migration-selection equilibrium without polymorphism at the T locus is reached rapidly following initial conditions (i.e., before mutation at the C locus occurs). Indeed, choosiness is neutral if there is no polymorphism at the T locus.

### Invasion of a choosier allele, and ESS choosiness

Our central issue, which is to study the evolution of choosiness which controls the strength of assortative mating, is tackled by assuming ancestral choosiness that allows the maintenance of polymorphism at the E and T loci and by numerically assessing the invasion conditions of choosiness alleles introduced in low frequency in a population at migration-selection equilibrium. We can thus determine the choosiness value, *α*_ESS_, that constitutes an evolutionary stable strategy (the ESS choosiness that is always favored) in cases where choosiness evolves alongside a magic trait, versus a pseudomagic trait complex, versus a nonmagic trait complex with physically unlinked loci.

More precisely, we start the population at the migration-selection equilibrium at the E and T loci given the ancestral choosiness, which is encoded by allele *C*_1_. We initially assume that allele *C*_1_ codes for the choosiness value *α*_1_ = *α*_poly_, the lowest value that allows the maintenance of polymorphism. The allele *C*_2_, coding for a higher choosiness value so that *α*_2_ = *α*_1_ + Δ*α*, is then introduced in linkage equilibrium with the other loci, in the same frequency (= 0.01) in both populations. After 200 generations, if the allele *C*_2_ has increased in frequency in the two populations we assume that it is able to invade and to completely replace the allele *C*_1_. We verified that invasion of the mutant allele at the C locus eventually leads to fixation, by checking that the mutant allele cannot be invaded by the allele coding for lower choosiness. If allele *C*_2_ is able to invade, we consider in another simulation that the ancestral allele *C*_1_ codes for the choosiness value previously encoded by allele *C*_2_, and we repeat the process by assessing whether an even choosier allele can invade. The evolutionarily stable choosiness *α*_ESS_ corresponds to the choosiness value encoded by *C*_1_ that is robust to invasion by a choosier allele *C*_2_. We checked that this allele coding for *α*_ESS_ is also robust to invasion by an allele coding for lower choosiness, so that *α*_2_ = *α*_1_ − Δ*α*. Δ*α* corresponds to the mutation effect size. To assess the ESS choosiness, we consider a small mutation effect size Δ*α* = 0.01, but in other analyses where we track the full invasion of a mutation, we consider larger mutation effect sizes.

### Contribution of linkage disequilibria to frequency change at the choosiness locus

The change in frequency at the C locus in population *k*, Δ*c*_2,*k*_, due to viability selection, mating and the production of zygotes can be decomposed into different components,

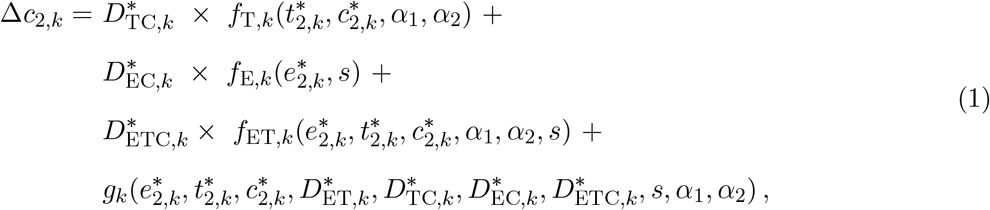

where *s* is the selection coefficient during viability selection, *α*_1_ and *α*_2_ are the choosiness values, and 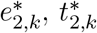, and 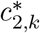 are the allelic frequencies after migration. Furthermore, 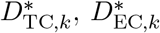, and 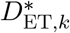 are the linkage disequilibria after migration between loci T and C, E and C, and E and T, respectively. Finally, 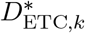 is the three-way linkage disequilibrium after migration between loci E, T and C.

The component 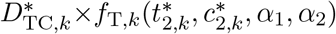 corresponds to the first-order contribution of 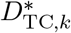 to the change in frequency at the C locus, with 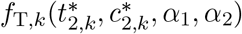 being the first-order approximation of the Barton-Turelli selection coefficient 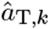 (this component can be isolated in this way because there is no direct selection on locus C, which could also contain terms to the first order in 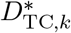 [24]). In other words, this term reflects how sexual selection at the locus T contributes to the evolution at the locus C via linkage disequilibrium. Likewise, the component 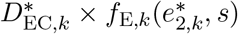 reflects how viability selection at the locus E contributes to the evolution at the locus C via linkage disequilibrium. The component 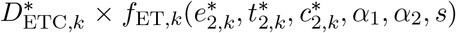 reflects how epistatic selection at the E and T loci contributes to the evolution at the locus C via three-way linkage disequilibrium. Finally, the term described by function *g* corresponds to the sum of all other higher-order contributions of 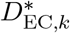, 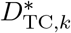 and 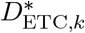, e.g., the terms associated with 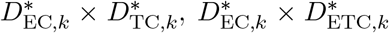, etc. This last term is generally negligible relative to the first three terms (Figs. S1 and S2).

We can show analytically that the functions *f*_T,*k*_ and *f*_ET,*k*_ are positive or negative depending on whether the local allelic frequencies at the E and T loci are greater or less than 1/2, and that the function *f*_E,*k*_ is negative for *k* = 1 and positive for *k* = 2 (see *Mathematica* notebook). Therefore, in our simulations where allelic divergence occurs (Fig. 1), the signs of the linkage disequilibria 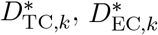, and 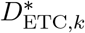 determine the direction of indirect sexual selection, indirect viability selection and indirect epistatic selection, respectively (Figs. S3 and S4); in particular, in population 2 where allelic frequencies at the E and T loci are greater than 1/2, positive (resp. negative) linkage disequilibria lead to positive (resp. negative) indirect selection on choosiness.

## RESULTS

### Magic traits, pseudomagic trait complexes, and the evolution of choosiness

With a magic trait (*r*_ET_ = 0), divergence at the pleiotropic locus, which is both under divergent ecological selection and used as a mating signal, and thus fulfills the functions of both E and T loci, is maximized for an intermediate choosiness value (black double-headed arrows in Fig. 1; as explained previously). In addition, the ESS choosiness promoted with a magic trait corresponds to the choosiness value that maximizes divergence (black single-headed arrows in Fig. 1). This ESS choosiness is promoted by indirect viability and sexual selection, through the build-up of linkage disequilibrium between the choosiness locus and the pleiotropic locus [16] (see also Supplementary Information Text). Because it is set by selection (in this case indirect), it is an example of partial reproductive isolation evolving as an adaptive optimum [25].

We now show that intermediate choosiness is favored not only with a magic trait (for *r*_ET_ = 0 in Fig. 2), but also with any nonmagic trait complex with separate ecological and mating signal loci (for *r*_ET_ *>* 0 in Fig. 2). We get qualitatively the same result if we consider the gene order CET instead of ETC (Fig. S5). Polymorphism at the mating signal locus T is maintained under random mating only with a magic trait (as shown for low choosiness in Fig. 1; see also Fig. S6). With a nonmagic trait complex, polymorphism at the T locus is lost under random mating, and the ESS choosiness can evolve only if at least a small level of choosiness is already present before secondary contact occurred (Figs. 1 and S6). If we condition on such initial maintenance of polymorphism, contrary to intuition, pseudomagic trait complexes characterized by physically linked loci (for 0 *< r*_ET_ *<* 0.5), and also to a lesser extent physically unlinked loci (for *r*_ET_ = 0.5), have the potential to favor the evolution of higher choosiness than do magic traits (Figs. 1 and 2). This effect is particularly pronounced under strong viability selection relative to migration, and when the choosiness modifier is linked to the other loci (*r*_TC_ *<* 0.5). Note that this effect can occur even if the choosiness modifier is unlinked (for *r*_TC_ = 0.5; this is not visible in Fig. 2, but see Fig. S7).

**Figure 2:**
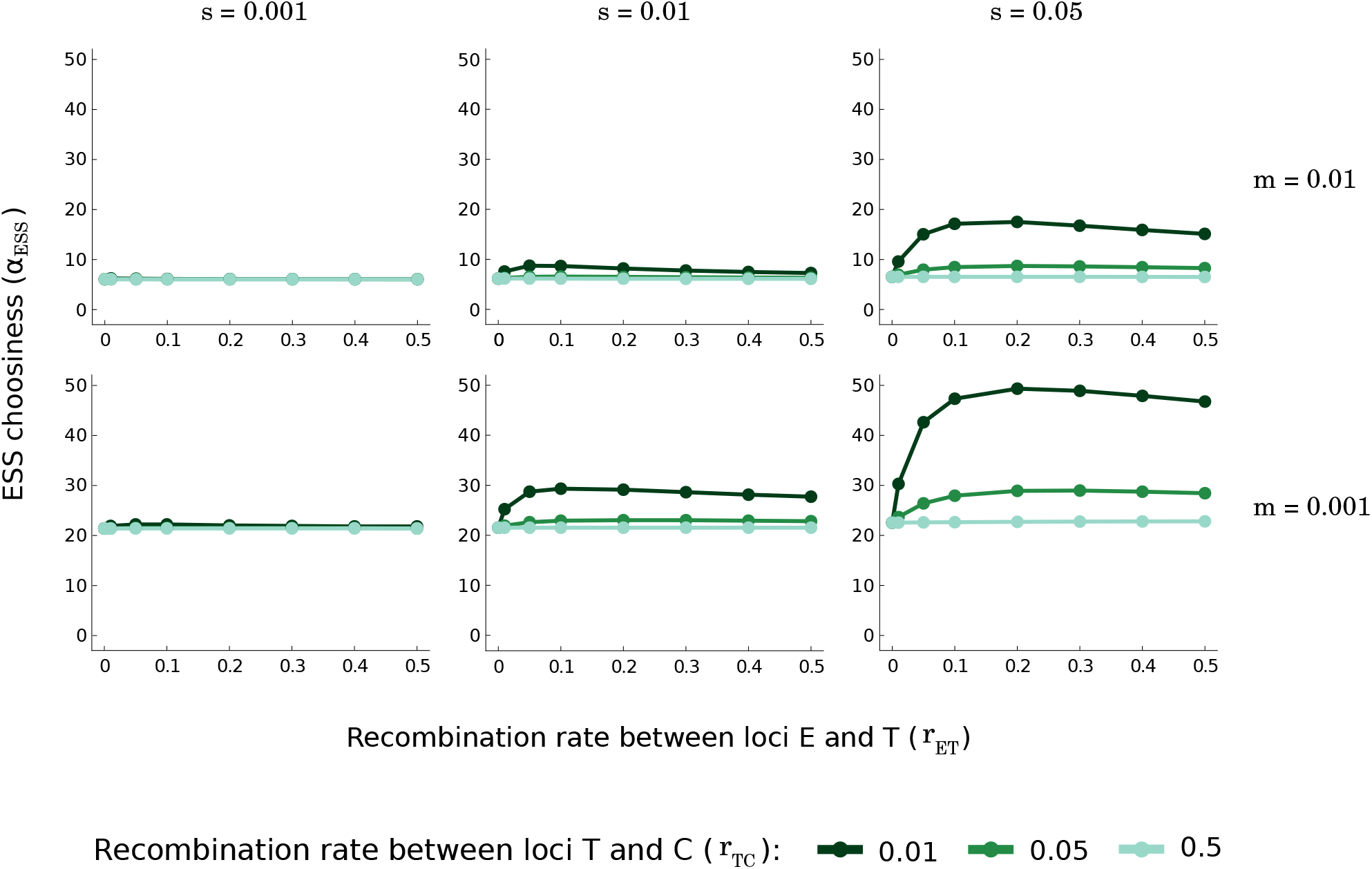
Choosiness at the evolutionary equilibrium depending on level of gene flow and genetic architecture. We represent the evolutionary stable choosiness, *α*_ESS_, depending on the selection coefficient (*s*), the migration rate (*m*), and the recombination rates (*r*_ET_ and *r*_TC_). The case of *r*_ET_ = 0 can be interpreted as a magic trait (remember that we assume that loci E and T are at maximum linkage disequilibrium, initially). For a given combination of parameters (*s, m*), an intermediate *r*_ET_ leads to the highest ESS choosiness; if *r*_ET_ is too small, three-way linkage disequilibrium and its effect on the evolution of high choosiness (as detailed in the main text) are too weak, and if *r*_ET_ is too high, recombination breaks linkage disequilibrium that causes indirect selection on choosiness (as detailed in Figs. S1 and S2). For *r*_TC_ = 0.5, changes in the recombination rate *r*_ET_ lead to slight changes in the ESS choosiness that are not visible here; in particular, for high ratio *s/m*, a nonmagic trait complex (*r*_ET_ *>* 0) can lead to a higher choosiness at evolutionary equilibrium than can be found with a magic trait (*r*_ET_ = 0) (see Fig. S7). The higher choosiness allowed by nonmagic trait complexes results in stronger reproductive isolation, as measured by a lower effective migration rate (see Fig. S8).

In our model, the higher choosiness that evolves alongside nonmagic trait complexes ultimately leads to stronger reproductive isolation than is seen with a magic trait, even though divergence at the T locus is lower (Fig. 1). We can show this by measuring the effective migration rate, which is inversely proportional to reproductive isolation (Fig. S8; note, however, that the measurement of reproductive isolation is still a matter of debate [26]).

Below, we will first describe the mechanism by which a nonmagic trait complex, and in particular a pseudomagic trait complex, can promote the evolution of higher choosiness than does a magic trait. We will then briefly explain why this mechanism is particularly efficient in promoting the evolution of choosiness when the choosiness modifier is tightly linked to the other loci (i.e., for a low *r*_TC_). Finally, we will outline the implication of a pseudomagic trait complex for the invasion of large-effect choosiness mutations.

### Mechanism by which a nonmagic trait complex promotes the evolution of high choosiness

Unlike with a magic trait, the ESS choosiness that is favored with a nonmagic trait complex does not maximize divergence at either the E or T locus (for *r*_ET_ *>* 0 in Fig. 1), which means that additional evolutionary forces come into play. In the case of a nonmagic trait complex, higher choosiness is favored because it prevents the deleterious consequences of recombination between the E and T loci, namely a mismatch between being locally adapted and having the locally preferred male phenotype. Such a higher choosiness is not favored in the case of a magic trait because recombination between the E and T loci is impossible by definition. As we now describe in detail, we can isolate this evolutionary force by dissecting the sources of indirect selection affecting the evolution of choosiness.

In the case of a nonmagic trait complex, indirect selection favoring the evolution of high choosiness relies on the build-up of an association among the choosiness, signal, and ecological loci. More precisely, this three-way linkage disequilibrium is characterized by a high frequency of choosier alleles, denoted *C*_2_ so that *α*_2_ *> α*_1_, being associated with the locally favored combination of alleles at the E and T loci (*E*_1_*T*_1_ in population 1, and *E*_2_*T*_2_ in population 2). If we artificially reduce this three-way linkage disequilibrium at the end of each generation over the course of simulations, the ESS choosiness with a nonmagic trait complex is no longer higher than the ESS choosiness with a magic trait (Figs. 3a and S9). This highlights the importance of three-way linkage disequilibrium for the evolution of higher choosiness in the case of a nonmagic trait complex.

**Figure 3:**
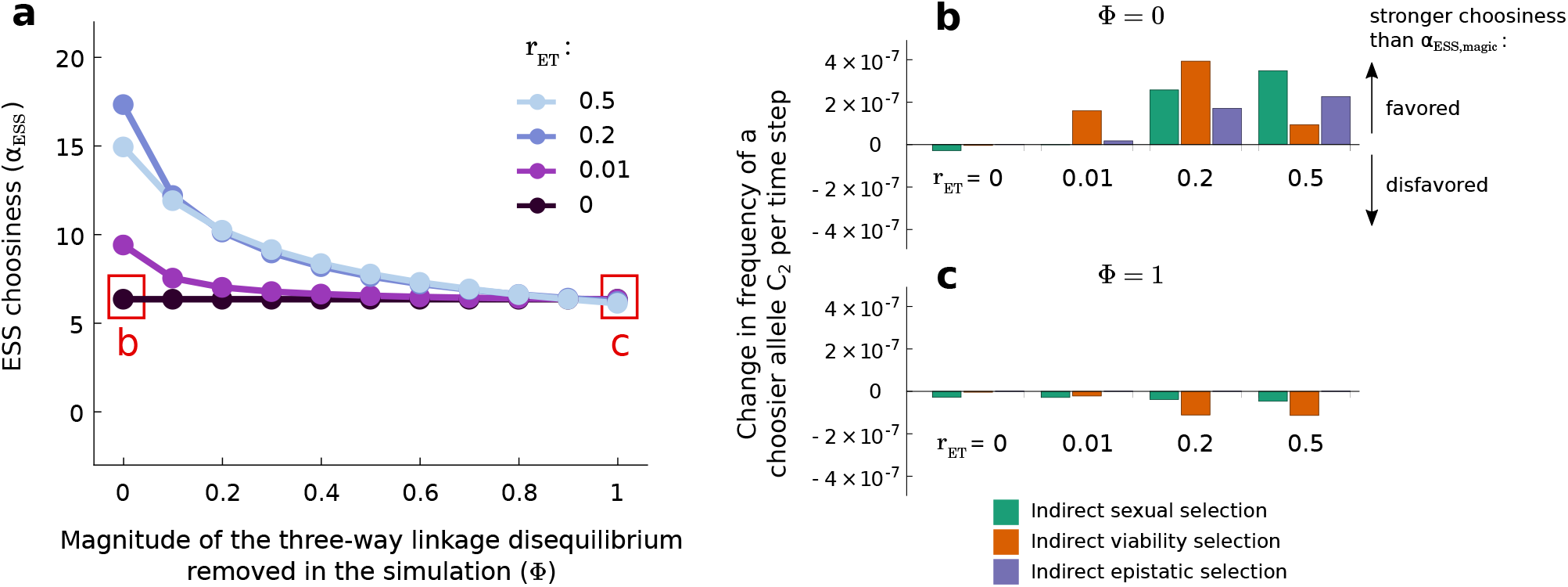
Choosiness at the evolutionary equilibrium depending on the magnitude of the three-way linkage disequilibrium that is removed artificially over the course of simulations. In panel **a**, we show the evolutionary stable choosiness, *α*_ESS_, depending on the recombination rate *r*_ET_ and the proportion of the three-way linkage disequilibrium, *D*_ETC_, removed artificially in the simulation. Over the course of the simulations, we reduce the magnitude of the three-way linkage disequilibrium by an amount that depends on the value Φ represented on the horizontal axis. At the end of each generation, we artificially reduce three-way linkage disequilibrium by transforming three-way linkage disequilibrium in each population *k* according to 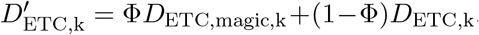, with *D*_ETC,magic,k_ = min [*D*_TC,k_(1 − 2*t*_2,*k*_), *D*_EC,k_(1 − 2*e*_2,*k*_)] being the equivalent measure to the three-way linkage disequilibrium if that formula were applied to the case of a magic trait. For Φ = 1, we thus artificially set the three-way linkage disequilibrium to its lowest possible value. In panels **b** and **c**, we represent the first-order contributions of indirect sexual selection (in green), indirect viability selection (in orange), and indirect epistatic selection (in purple), to the evolution of stronger choosiness than the ESS choosiness value favored in the case of a magic trait, *α*_ESS,magic_, depending on whether the three-way linkage disequilibrium is reduced to its lowest possible value over the course of simulations (**c**) or not (**b**). Choosiness *α*_1_ is set to be the ESS choosiness value obtained for *r*_ET_ = 0 (*α*_1_ = *α*_ESS,magic_). For each combination of parameters, we know if a choosier allele *C*_2_, coding for *α*_2_ = *α*_1_ + 1, will increase or decrease in frequency based on the ESS value we were able to determine in the analysis done in panel **a**. We therefore implement *C*_2_ at a frequency equal to 0.01 if it is destined to increase in frequency, or equal to 0.99 if it is destined to decrease in frequency. Over the course of the simulation, we measure the mean first-order contributions of linkage disequilibria to Δ*c*_2,2_, corresponding to the first three terms of Equation 1, while the frequency of the choosier allele *c*_2,2_ is between 0.05 and 0.95. These first order-contributions of linkage disequilibria correspond to first-order approximations of the effect of indirect sexual selection, indirect viability selection, and indirect epistatic selection on the change in frequency of the choosier allele. See Figs. S1 and S2 for more details on the effect of *r*_ET_ on the contributions of selection to the evolution of choosiness, shown in panel **b**. Here, *m* = 0.01, *s* = 0.05 and *r*_TC_ = 0.01. In panels **b** and **c**, *α*_ESS,magic_ = 6.54 (estimated numerically; see position of the red squares in panel **a**).

Three-way linkage disequilibrium is established because, under secondary contact, higher choosiness reduces the risk of producing recombinant offspring. A choosier allele *C*_2_ is more likely to remain associated with the locally favored allelic combination at the E and T loci than is a less choosy allele *C*_1_. For instance, in population 2, *E*_2_*T*_2_*C*_2_ females mate less often with maladapted *E*_1_*T*_1_ males than do less choosy *E*_2_*T*_2_*C*_1_ females, and thus produce recombinant offspring *E*_2_*T*_1_ and *E*_1_*T*_2_ less often. This results in the maintenance of the beneficial allelic association *E*_2_*T*_2_ with *C*_2_ rather than with *C*_1_, leading to the build-up of three-way linkage disequilibrium in the case of a nonmagic trait complex (Fig. S10). This contrasts with the case of a magic trait where there is no recombination between the E and T loci (*r*_ET_ = 0) and thus the build-up of an association akin to this three-way linkage disequilibrium cannot occur. Note that, with a magic trait, the formula for three-way linkage disequilibrium can still be applied (e.g., to measure selection for *r*_ET_ = 0 in Fig. 3), but results in a measure proportional to two-way linkage disequilibrium [24] (see also *Mathematica* notebook).

The establishment of three-way linkage disequilibrium that occurs only alongside nonmagic trait complexes contributes to indirect epistatic selection (non-additive fitnesses) favoring the evolution of higher choosiness, beyond the ESS choosiness promoted with a magic trait (purple bars in Fig. 3b, and see also time series in Fig. S3). In other words, with a nonmagic trait complex, viability and sexual selection act non-additively on choosiness via the association between the choosier allele *C*_2_ and the locally favored combination of alleles at the E and T loci (*E*_1_*T*_1_ in population 1 and *E*_2_*T*_2_ in population 2). Obviously, if we artificially reduce the three-way linkage disequilibrium, indirect epistatic selection no longer favors the evolution of high choosiness (almost vanishing purple bars in Fig. 3c).

Additionally, three-way linkage disequilibrium contributes to the build-up of two-way linkage disequilibria between loci T and C during viability selection, and between loci E and C during sexual selection (see Supplementary Information Text). These two-way linkage disequilibria then lead to indirect sexual and viability selection favoring the evolution of higher choosiness than the ESS choosiness established with a magic trait (green and orange bars in Fig. 3b, and see also time series in Figs. S3 and S4). Indeed, if we artificially reduce the three-way linkage disequilibrium, indirect sexual and viability selection no longer favor the evolution of choosiness higher than with a magic trait (green and orange bars in Fig. 3c); this is because three-way linkage disequilibrium no longer leads to the build-up of positive two-way linkage disequilibria.

In supplementary analyses, we find that the difference in ESS choosiness evolving alongside a nonmagic and a magic trait can even be amplified by a weak cost of choosiness (Supplementary Information text; Fig. S11). We also show that in a diploid version of the model, nonmagic trait complexes can similarly promote the evolution of stronger assortative mate choice than do magic traits (Supplementary Information text; Figs. S12 and S13).

To summarize, a nonmagic trait complex allows the establishment of an association among the choosiness, signal, and ecological loci (three-way linkage disequilibrium). With few exceptions, detailed below, this association favors the evolution of higher choosiness than the ESS choosiness established with a magic trait (Figs. 2 and S7). In other words, with a nonmagic trait complex, this higher choosiness is favored because it prevents the deleterious consequences of recombination between signal and ecological loci. This recombination does not occur in the case of a magic trait.

### A pseudomagic trait complex can promote the evolution of higher choosiness than do unlinked loci

Although three-way linkage disequilibrium may ultimately favor the evolution of choosiness alongside any nonmagic trait complex, a pseudomagic trait complex (*r*_ET_ *<* 0.5) can promote the evolution of higher choosiness than do physically unlinked loci (*r*_ET_ = 0.5; Fig. 2). This is because with the gene order ETC, recombination between the E and T loci degrades the linkage disequilibrium between the E and C loci and thus may decrease the contribution of indirect viability selection to the evolution of choosiness (orange bars for *r*_ET_ = 0.2 vs. 0.5 in Fig. 3b; see also Figs. S1 and S2; intuitively, this is related to the advantage of a magic trait in preventing recombination). Likewise, with the gene order CET, recombination between the E and T loci degrades the linkage disequilibrium between the T and C loci and thus may decrease the contribution of indirect sexual selection to the evolution of choosiness. As a result, indirect selection favoring the evolution of high choosiness is the strongest alongside a pseudomagic trait complex (Fig. 2).

### Tight linkage between the choosiness and pseudomagic trait loci favors higher choosiness

As noted above, the degree to which nonmagic trait complexes (*r*_ET_ *>* 0) favor the evolution of higher choosiness than do magic traits (*r*_ET_ = 0) depends on the recombination rate, *r*_TC_, between the choosiness and the trait loci. This effect is especially pronounced when the choosiness modifier itself is tightly linked to the other loci (e.g., for *r*_TC_ = 0.01 in Fig. 2; or for *r*_CE_ = 0.01 in Fig. S5 where we consider the gene order CET instead of ETC). This is because low recombination between the choosiness modifier locus and the other loci reduces the indirect selection that inhibits the evolution of very high choosiness and that therefore leads to intermediate choosiness [16] (see also Supplementary Information Text).

In contrast, when the choosiness modifier is loosely linked to the trait loci, a nonmagic trait complex may even lead to slightly lower choosiness than the ESS choosiness established with a magic trait (e.g., for *r*_TC_ = 0.5 but the effect is so small that this is not visible in Fig. 2; see Fig. S7). This is because recombination between the E and T loci degrades the linkage disequilibrium that causes indirect viability selection on choosiness (as detailed in Figs. S1 and S2).

### Pseudomagic trait complexes and the invasion of large-effect choosiness mutations

Now that we have explained why the choosiness at evolutionary equilibrium can be higher for a nonmagic trait complex, and in particular for a pseudomagic trait complex, than for a magic trait, we turn to the investigation of the implication of nonmagic trait complexes for the spread or loss of large-effect choosiness mutations (Fig. 4; see Fig. S14 for a log-scale highlighting the slight increase or the decrease in frequency of these mutations). We track the change in frequency of choosier alleles for different mutation effects, when ancestral choosiness in the population is low, but high enough to maintain polymorphism at the E and T loci.

**Figure 4:**
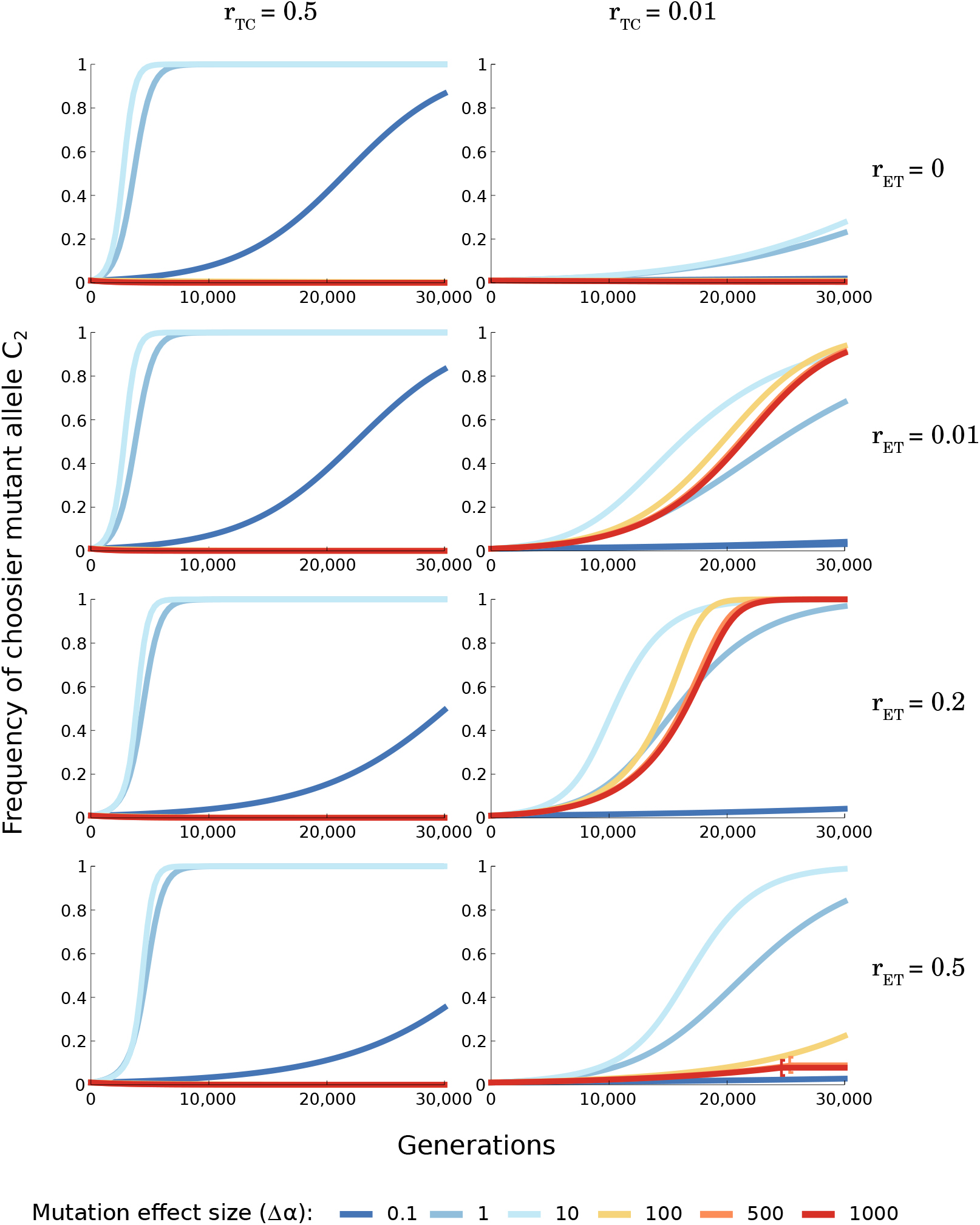
Time series of the invasion of mutant alleles at the choosiness locus for different mutation effect sizes and different genetic architectures. Choosiness *α*_1_ is set to be the lowest choosiness value that maintains polymorphism at the T locus for all recombination rates tested (*α*_1_ = 0.13; estimated numerically). We then show the invasion in population 2 of a mutant allele coding for a choosiness *α*_2_ = *α*_1_ + Δ*α*. We observe the same invasion dynamics in population 1 (not shown). Brackets show the time points where choosiness becomes a neutral trait because polymorphism at the T locus is lost (bottom right panel). Here, *m* = 0.01 and *s* = 0.05.

For *r*_TC_ = 0.5, mutant alleles that code for higher choosiness, but that are not overly choosy, can invade (Fig. 4). The recombination rate between loci encoding the ecological trait and the mating signal (*r*_ET_) has little effect on the dynamics of invasion, unless the mutation effect size is very small. This is not surprising given that the ESS choosiness depends very little on *r*_ET_ (Fig. S7).

For *r*_TC_ = 0.01, a nonmagic trait complex (*r*_ET_ *>* 0; especially pseudomagic trait complexes with intermediate *r*_ET_) can favor the invasion of choosier mutant alleles than does a magic trait (*r*_ET_ = 0) (Fig. 4). This is in line with the high ESS choosiness obtained with a nonmagic trait complex (Fig. 2). Interestingly, a pseudomagic trait complex allows the evolution of very high choosiness through the spread of a single large-effect mutation (so that Δ*α* ≥ 100 in Fig. 4) which far overshoots the choosiness value at evolutionary equilibrium (see also pairwise invasibility plots in Fig. S15).

With a nonmagic trait complex, the invasion of choosier mutant alleles occurs only if choosiness is initially strong enough to maintain polymorphism. If mating is initially random, the quick loss of polymorphism after secondary contact, especially when loci are physically unlinked, impedes the invasion of any choosier alleles (Fig. S16; in contrast, with a magic trait, polymorphism is always maintained, even under random mating). Assuming that polymorphism at the T locus is initially maintained, we get qualitatively the same invasion time series as in the symmetric case (in Fig. 4) if migration is asymmetric (Fig. S17), viability selection is asymmetric (Fig. S18), or choosiness is asymmetric (Fig. S19). Such asymmetric conditions cause the loss of polymorphism at the mating signal locus to occur more easily than under symmetric conditions (Fig. S20). If polymorphism is maintained at the outset, however, pseudomagic trait complexes predominantly favor the evolution of higher choosiness than does a magic trait.

## DISCUSSION

Our model shows the fragility of the intuitive prediction that magic traits, subject to divergent selection and pleiotropically affecting reproductive isolation, necessarily favor stronger reproductive isolation than other genetic architectures. We show that, although magic traits favor divergence and the maintenance of polymorphism, pseudomagic trait complexes, characterized by separate physically linked loci being subject to divergent selection and affecting reproductive isolation, have the potential to promote the evolution of stronger assortative mate choice than do magic traits, provided that polymorphism at the ecological and mating signal loci is maintained. In the case of a pseudomagic trait complex, this strong assortative mate choice is favored because it ultimately diminishes the risk of recombining the genes that underlie favorable complexes of ecological and mating traits. It thus reduces the production of recombinant offspring with lower average fitness than the parental forms. This recombination risk does not exist in the case of magic traits (which rely on pleiotropy, by definition), explaining why this strong assortative mate choice is favored in the case of pseudomagic trait complexes and in the case of nonmagic trait complexes with physically unlinked loci (although in the latter case polymorphism is particularly difficult to maintain, and the spread of a single large-effect choosiness mutation can only slightly overshoot the ESS choosiness value).

With a magic trait, a single pleiotropic locus is both under ecologically divergent selection and involved in assortative mating [11]. Functionally, this is analogous to a situation where linkage disequilibrium between loci controlling ecological and mating traits is maximized and cannot be broken by recombination. Nevertheless, one should not make the mistake of equating linkage disequilibrium between a subset of loci with increased reproductive isolation [7]. Clearly, incipient species are characterized not only by genes involved in premating isolation but also by other genes, such as neutral genes and genes involved in postzygotic isolation, and one must consider the linkage disequilibrium among all of these loci to infer the likelihood of speciation. Importantly, the extent to which linkage disequilibrium between all genes is maintained depends on the gene flow between incipient species and thus on the strength of assortative mate choice [23]. Therefore, whether a given genetic architecture favors speciation strongly depends on the strength of assortative mate choice that can evolve.

In a previous study [23], the authors concluded that it is not necessarily important to identify whether co-localizing mating and ecological components are pleiotropic or merely tightly linked, in terms of evolutionary divergence. By assessing the impact of genetic linkage on the build-up of premating isolation via the evolution of strong assortative mate choice, our study shows that a very different conclusion applies to reproductive isolation more broadly. We predict that a pseudomagic trait complex characterized by very tight linkage (e.g., for *r*_ET_ ∈ [0.01, 0.1]) is a genetic architecture that is prone to favor the evolution of strong reproductive isolation, provided that polymorphism at the ecological and mating signal loci is maintained after secondary contact. Although a magic trait is more likely to maintain polymorphism in this situation (especially under random mating), the strength of assortative mate choice that can evolve with such a magic trait is limited. In contrast, a pseudomagic trait complex more easily leads to the loss of polymorphism at the mating signal locus upon secondary contact, but if it does not, then it can lead to the evolution of stronger assortative mate choice than does a magic trait. In this respect, a pseudomagic trait complex is more likely to maintain species-specific allelic combinations, and may be more effective in the promoting later stages of speciation than a magic trait.

The recombination rate between loci involved in premating isolation can evolve upon secondary contact, which we did not consider in our model. For example, chromosomal rearrangements (e.g., inversions) have suppressed recombination between loci that may be involved in premating isolation in some taxa [27–29], as predicted by population genetics theory [30]. Our study highlights that this local suppression of recombination may not necessarily favor the establishment of strong premating isolation, but may instead inhibit the evolution of mate choice that leads to reduced gene flow across the genome.

Our model also emphasizes the importance of the location in the genome of genetic loci encoding choosiness, i.e., encoding the strength of assortative mate choice. Under secondary contact, pseudomagic trait complexes favor the evolution of strong assortative mate choice only when choosiness loci are tightly linked to loci forming pseudomagic trait complexes. This raises the question of whether the evolution of high choosiness could be limited by the genetic architecture and position of loci encoding choosiness. For instance, if choosiness is a quantitative trait encoded by loci distributed uniformly along the genome, then the evolution of choosiness is likely to be limited by number of choosiness loci that are tightly linked to ecological and mating signal loci, rather than by indirect selection favoring choosiness. Such a prediction calls for more empirical investigation of the genetic basis of choosiness, which has received little attention so far. Indeed, intraspecific variation in choosiness is required to assess quantitative trait loci along genomes. Because models predict that choosiness alleles may spread uniformly through incipient species during speciation with gene flow (because choosiness operates through a “one-allele” mechanism [7]), there may be limitations to the dissection of the genetic basis of choosiness if most alleles go to fixation. However, by measuring preference behaviors in hybrids that may show variation in choosiness this limitation could conceivably be overcome (as in [28] in the case of the preference for conspecific versus heterospecific mates), shedding light on the constraints to the evolution of choosiness alongside pseudomagic trait complexes. Before the genetic architecture of choosiness can be assessed, however, it is necessary to develop a standardized measure of the choosiness [31], as a stronger preference can be independent of choosiness when there is variation in the deviation between the preference and the trait values of the individuals.

We appreciate that our modelling approach comes with its limitations, including a limited number of loci, a geographical context of secondary contact, and a specific mating rule (phenotype matching). As a whole, however, our model highlights the importance of genetic architecture for the evolution of assortative mating and thus the build-up of premating isolation. Speciation with gene flow has been traditionally studied under the assumption that ‘magic’ genes encode traits involved in premating isolation [8, 13–17], and some authors have speculated that nonmagic trait complexes could mimic the role of magic traits in the speciation process [18, 21, 22]. Our study shows that by reducing the risk of recombination, strong assortative mate choice can evolve alongside a nonmagic trait complex, and even more so alongside a pseudomagic trait complex, thus promoting the build-up of stronger premating isolation than would occur alongside a magic trait. We hope that the predictions of our model will stimulate further empirical investigations assessing the genetic architecture underlying premating isolation.

## ACKNOWLEDGMENTS

We thank B. Lerch, K. Xu and two anonymous reviewers for comments that have helped improve our manuscript.

## AUTHORSHIP STATEMENT

All authors conceived the idea, TGA developped the model and performed analyses, TGA wrote the first draft of the manuscript, and all authors contributed substantially to revisions.

## DATA ACCESSIBILITY STATEMENT

The *Mathematica* notebook for this paper is archived on Zenodo: https://doi.org/10.5281/zenodo.7631806

