## Supplementary Information for "The effectiveness of pseudomagic traits in promoting premating isolation"

#### Contents

**Pages 2 - 12: Supplementary Information Text**

- Pages 2 - 3 Main recursion equations
- Pages 4 - 6 Recursion equations during viability selection and sexual selection
- Page 6 With a magic trait, the ESS choosiness maximizes divergence at the pleiotropic locus
- Page 7 Low recombination between the choosiness modifier and other loci reduces the constraint on the evolution of very high choosiness
- Pages 7 - 10 With weak costs of choosiness, pseudomagic trait complexes promote the evolution of higher choosiness than magic traits
- Pages 10 - 12 With a diploid life cycle, pseudomagic trait complexes can promote the evolution of higher choosiness than magic traits

**Pages 13 - 34: Figures S1 to S22**

### Supplementary Information Text

#### Main recursion equations

##### (a) Genotypes

Following Servedio and Bürger (2020, *Evolution* **74**(11):2438-2450), we consider a two-island population genetics model of haploids with assortative mating that occurs by phenotype matching. We follow changes in allelic frequencies at three autosomal diallelic loci, one coding for an ecological trait, E, one coding for a trait used as a mating signal (or "mating trait"), T, and one coding for choosiness during mate choice, C. The ecological locus E is subject to divergent viability selection, such that $E_1$ is locally adapted in population 1 and $E_2$ is locally adapted in population 2. The signal locus T is subject to sexual selection, such that females with allele $T_1$ or $T_2$ prefer to mate with males with a matching trait allele, regardless of which population they are in, thereby leading to variation in mating success in males. In our model, $T_1$ is generally characteristic of population 1 and $T_2$ is generally characteristic of population 2, generating divergent sexual selection at the T locus. The choosiness locus C controls the strength of preference, such that females mate preferentially with matching males by a factor controlled by the allele at the C locus. In particular, we assume that females with allele $C_2$ are choosier than those with allele $C_1$ (note that subscripts 1 and 2 describing alleles at the C locus do not correspond to the habitat in which alleles at the E and T loci usually predominate). We assume that the gene order is ETC, so that recombination occurs at a rate $r_{\text{ET}}$ between loci E and T, and $r_{\text{TC}}$ between loci T and C. We also assume there is no crossover interference, so that the rate of recombination between loci E and C is $(1 - r_{\text{ET}})\, r_{\text{TC}} + r_{\text{ET}}\, (1 - r_{\text{TC}})$.

There are thus eight genotypes, $E_1T_1C_1$, $E_2T_1C_1$, $E_1T_2C_1$, $E_2T_2C_2$, $E_1T_1C_2$, $E_2T_1C_2$, $E_1T_2C_2$, and $E_2T_2C_2$ the frequencies of which are denoted by $x_{1,k}$, through $x_{8,k}$, respectively, where the second subscript $k$ denotes the population (either 1 or 2). We also follow frequencies $e_{2,k}$, $t_{2,k}$, and $c_{2,k}$ of the alleles $E_2$, $T_2$ and $C_2$ alleles, respectively (again, the subscript $k$ denotes the population). The life cycle is as follow: census, migration, viability selection, mating, production of zygotes.

##### (b) Migration

After migration, the frequencies of the genotypes are denoted with a superscripted asterisk, such that $x^*_{i,k} = (1-m_k)\, x_{i,k} + m_k\, x_{i,l}$, where $l = 2$ when $k = 1$ and $l = 1$ when $k = 2$. Parameter $m_k$ denotes the proportion of the population $k$ that consists of migrants after migration has occurred. We consider both symmetric and asymmetric migration, but unless we specify otherwise we assume symmetric migration such that $m_1 = m_2 = m$.

##### (c) Viability selection

Divergent viability selection acts on the ecological locus in both sexes. Individuals with allele $E_k$ at the E locus have a selective advantage due to local adaptation, with relative fitness $1 + s_k$ in population

$k$. The frequencies of the genotypes after viability selection are denoted with a superscripted double asterisk, such that:

$$x_{i,k}^{**} = \frac{(1 + d_{ik} s_k)\, x_{i,k}^{*}}{1 + s_k\, e_{k,k}^{*}}\,. \tag{S1}$$

Here, $d_{ik} = 1$ if genotype $i$ has allele $E_k$ and $d_{ik} = 0$ otherwise, and $e_{k,k}^{*}$ represents the frequency of allele $E_k$ (first subscript) in population $k$ (second subscript) after migration (thus, $e_{1,1}^{*} = \sum_{i \in \{1,3,5,7\}} x_{i,1}^{*}$ and $e_{2,2}^{*} = \sum_{i \in \{2,4,6,8\}} x_{i,2}^{*}$). We consider both symmetric and asymmetric viability selection, but unless we specify otherwise we assume symmetric viability selection such that $s_1 = s_2 = s$.

#### (d) Mating

Nonrandom mating occurs by phenotype matching. Specifically, females with alleles $C_n$ and carrying allele $T_1$ (resp. $T_2$) are $1 + \alpha_n$ times as likely to mate with a $T_1$ male (resp. a $T_2$ male) than with a male of the opposite $T$ allele, if she encounters one of each. We assume that mating preference coefficients are the same in the two populations. In population $k$, the frequency of mated pairs of females with genotype $i$ and males with genotype $j$ is thus:

$$M_{ij,k} = \frac{(1 + \delta_{c_1,ij}\alpha_1)\,(1 + \delta_{c_2,ij}\alpha_2)\, x_{j,k}^{**}\, x_{i,k}^{**}}{\sum_z (1 + \delta_{c_1,iz}\alpha_1)\,(1 + \delta_{c_2,iz}\alpha_2)\, x_{z,k}^{**}}\,. \tag{S2}$$

where $\delta_{c_1,ij} = 1$ (resp. $\delta_{c_2,ij} = 1$) if the female genotype $i$ carries allele $C_1$ (resp. $C_2$) at the choosiness locus and male genotype $j$ carries the same allele as the female genotype $i$ at the trait locus, and $\delta_{c_1,ij} = 0$ (resp. $\delta_{c_2,ij} = 0$) otherwise. The normalization in the denominator ensures that females have equal mating success regardless of their genotype. We refer to the mating preference coefficients, $\alpha_1$ and $\alpha_2$, as the 'choosiness values' encoded by alleles $C_1$ and $C_2$. We consider both symmetric choosiness, such that $T_1$ and $T_2$ females have the same choosiness, and asymmetric choosiness, but unless we specify otherwise we assume symmetric choosiness.

#### (e) Production of zygotes

Mating brings two haploid genomes together and is followed immediately by meiosis. During meiosis, recombination occurs at a rate $r_{\mathrm{ET}}$ between loci E and T, and $r_{\mathrm{TC}}$ between loci T and C. In each population, the expected genotype frequencies of zygotes in the next generation are therefore calculated by summing the appropriate mating frequencies $M_{ij,k}$ assuming Mendelian segregation and recombination in haploids.

This sequence of equations leads to the genotypic frequencies after reproduction in terms of the genotypic frequencies $x_{i,k}$ before migration, assuming that generations do not overlap. These genotypic recursion equations can be transformed to calculate frequencies of the alleles and the linkage disequilibria between the E, T, and C loci.

#### Recursion equations during viability selection and sexual selection

Here, we focus on the changes in frequencies and linkage disequilibria in population 2. Analogous equations apply in population 1.

##### (a) Viability selection

After migration:

- $e_2^*$ represents the frequency of allele $E_2$
- $t_2^*$ represents the frequency of allele $T_2$
- $c_2^*$ represents the frequency of allele $C_2$
- $D_{\mathrm{ET}}^*$ represents the linkage disequilibrium between loci E and T
- $D_{\mathrm{TC}}^*$ represents the linkage disequilibrium between loci T and C
- $D_{\mathrm{EC}}^*$ represents the linkage disequilibrium between loci E and C
- $D_{\mathrm{ETC}}^*$ represents the three-way linkage disequilibrium between loci E, T and C

Allelic frequencies change during viability selection (superscript 'vs' stands for viability selection) following

$$\Delta^{\mathrm{vs}}\, e_2 = (1 - e_2^*)\, e_2^*\, \hat{a}_{\mathrm{E}}^* \tag{S3}$$

$$\Delta^{\mathrm{vs}}\, t_2 = D_{\mathrm{ET}}^*\, \hat{a}_{\mathrm{E}}^* \tag{S4}$$

$$\Delta^{\mathrm{vs}}\, c_2 = D_{\mathrm{EC}}^*\, \hat{a}_{\mathrm{E}}^*\,, \tag{S5}$$

where

$$\hat{a}_{\mathrm{E}}^* = \frac{s_2}{1 + e_2^* s_2} \tag{S6}$$

and corresponds to the selection coefficient during viability selection.

Changes in two-way linkage disequilibria during viability selection can be expressed as

$$\Delta^{\mathrm{vs}}\, D_{\mathrm{ET}} = \hat{a}_{\mathrm{E}}^*\, D_{\mathrm{ET}}^*\, h^* \tag{S7}$$

$$\Delta^{\mathrm{vs}}\, D_{\mathrm{TC}} = \hat{a}_{\mathrm{E}}^*\, \left(D_{\mathrm{ETC}}^* - D_{\mathrm{EC}}^*\, D_{\mathrm{ET}}^*\, \hat{a}_{\mathrm{E}}^*\right) \tag{S8}$$

$$\Delta^{\mathrm{vs}}\, D_{\mathrm{EC}} = \hat{a}_{\mathrm{E}}^*\, D_{\mathrm{EC}}^*\, h^*\,, \tag{S9}$$

where

$$h^* = \frac{1 - e_2^*\,(2 + e_2^* s_2)}{1 + e_2^* s_2}\,. \tag{S10}$$

The expression of the change in three-way linkage disequilibrium, $\Delta^{\mathrm{vs}}\, D_{\mathrm{ETC}}$, is unwieldy and can be found in the *Mathematica* notebook.

Factoring by $\hat{a}_{\mathrm{E}}^{*} > 0$, as we did, simplifies the interpretation of these expressions. In particular, it is clear that increased three-way linkage disequilibrium, $D_{\mathrm{ETC}}^{*}$, leads to the build-up of two-way linkage disequilibrium between loci T and C: $\Delta^{\mathrm{vs}}\, D_{\mathrm{TC}}$ increases as $D_{\mathrm{ETC}}^{*}$ increases.

#### (b) Sexual selection

After viability selection:

- $e_2^{**}$ represents the frequency of allele $E_2$
- $t_2^{**}$ represents the frequency of allele $T_2$
- $c_2^{**}$ represents the frequency of allele $C_2$
- $D_{\mathrm{ET}}^{**}$ represents the linkage disequilibrium between loci E and T
- $D_{\mathrm{TC}}^{**}$ represents the linkage disequilibrium between loci T and C
- $D_{\mathrm{EC}}^{**}$ represents the linkage disequilibrium between loci E and C
- $D_{\mathrm{ETC}}^{**}$ represents the three-way linkage disequilibrium between loci E, T and C

We then consider the genotypic frequencies of mating males, allowing us to express changes in allelic frequencies and linkage disequilibria due to sexual selection alone.

Allelic frequencies change during sexual selection (superscript 'ss' stands for sexual selection) following

$$\Delta^{\mathrm{ss}}\, e_2 = D_{\mathrm{ET}}^{**}\, \hat{a}_{\mathrm{T}}^{**} \tag{S11}$$

$$\Delta^{\mathrm{ss}}\, t_2 = (1 - t_2^{**})\, t_2^{**}\, \hat{a}_{\mathrm{T}}^{**} \tag{S12}$$

$$\Delta^{\mathrm{ss}}\, c_2 = D_{\mathrm{TC}}^{**}\, \hat{a}_{\mathrm{T}}^{**}\,, \tag{S13}$$

where $\hat{a}_{\mathrm{T}}^{**}$ is a function of $\alpha_1$, $\alpha_2$, $c_2^{**}$, $t_2^{**}$, and $D_{\mathrm{TC}}^{**}$, and corresponds to the selection coefficient during sexual selection. Its expression is unwieldy and given in the *Mathematica* notebook. When sexual selection favors the allele $T_2$ in population 2, as it is the case in our simulations, we have $\hat{a}_{\mathrm{T}}^{**} > 0$.

Changes in two-way linkage disequilibria during sexual selection can be expressed as

$$\Delta^{\mathrm{ss}}\, D_{\mathrm{ET}} = \hat{a}_{\mathrm{T}}^{**}\, D_{\mathrm{ET}}^{*}\, h^{**} \tag{S14}$$

$$\Delta^{\mathrm{ss}}\, D_{\mathrm{TC}} = \hat{a}_{\mathrm{T}}^{**}\, D_{\mathrm{TC}}^{*}\, h^{**} \tag{S15}$$

$$\Delta^{\mathrm{ss}}\, D_{\mathrm{EC}} = \hat{a}_{\mathrm{T}}^{**}\, \left(D_{\mathrm{ETC}}^{**} - D_{\mathrm{TC}}^{**}\, D_{\mathrm{ET}}^{**}\, \hat{a}_{\mathrm{T}}^{**}\right)\,, \tag{S16}$$

where $h^{**}$ is a function of $\alpha_1$, $\alpha_2$, $c_2^{**}$, $t_2^{**}$, and $D_{\mathrm{TC}}^{**}$. It is given in the *Mathematica* notebook because its expression is unwieldy.

The expression of the change in three-way linkage disequilibrium, $\Delta^{\mathrm{ss}}\, D_{\mathrm{ETC}}$, is unwieldy and can be found in the *Mathematica* notebook.

Factoring by $\hat{a}_{\mathrm{T}}^{**} > 0$, as we did, simplifies the interpretation of these expressions. In particular, it is clear that increased three-way linkage disequilibrium, $D_{\mathrm{ETC}}^{*}$, leads to the build-up of two-way linkage disequilibrium between loci E and C: $\Delta^{\mathrm{ss}}\ D_{\mathrm{EC}}$ increases as $D_{\mathrm{ETC}}^{*}$ increases.

#### With a magic trait, the ESS choosiness maximizes divergence at the pleiotropic locus

With a magic trait, there is a pleiotropic locus that fulfills the functions of both the E and the T loci, because it is under divergent ecological selection and used as a mating signal (for simplicity, we refer to it in this section as locus T). Divergence at this locus is maximized for an intermediate choosiness value (Fig. 1a-b for $r_{\mathrm{ET}} = 0$ assuming maximum linkage disequilibrium initially; see also Servedio 2011, *Proc. Biol. Sci.* **278**(1703):179-187). Indeed, very high choosiness causes rare, very choosy females to mate with rare males in proportion to their frequency, resulting in the loss of positive frequency-dependent sexual selection and therefore lower divergence in the mating signal. Indirect viability and sexual selection favor the evolution of an intermediate level of choosiness that maximizes divergence through the build-up of linkage disequilibrium between the choosiness and mating signal loci (Servedio, 2011). As indicated by the fact that intermediate choosiness maximizes divergence at the T locus, intermediate choosiness increases the production of offspring carrying the locally favored allele at the T locus. As a result, an allele coding for such divergence-maximizing choosiness at the C locus is more strongly associated with the locally favored allele at the T locus than is any other choosiness allele, including those coding for very high choosiness. This leads to the build-up of positive linkage disequilibrium between loci T and C.

It is somewhat counter-intuitive that an allele coding for higher choosiness is not necessarily more likely to be associated with the locally favored allele at the T locus. Although such a very choosy allele reduces the proportion of crosses that break its association with the beneficial allele at the locus T via recombination (e.g., crosses between $T_2C_2$ and $T_1C_1$ individuals in population 2), it also reduces the proportion of crosses that build an association with the beneficial allele at the T locus via recombination (e.g., crosses between $T_1C_2$ and $T_2C_1$ individuals in population 2) (Fig. S21). Due to this latter effect of high choosiness, an allele coding for intermediate choosiness that is closer to the choosiness that maximizes divergence is more strongly associated with the locally favored allele at the T locus; a similar effect would occur if intermediate choosiness evolve from high choosiness.

#### Low recombination between the choosiness modifier and other loci reduces the constraint on the evolution of very high choosiness

In the previous subsection, we explained why the ESS choosiness that maximizes divergence at the T locus is intermediate with a magic trait (as in the previous subsection, T here refers to the pleiotropic locus, for simplicity). Importantly, the build-up of linkage disequilibrium favoring the evolution of the ESS choosiness relies on the recombination between loci T and C, $r_{\mathrm{TC}}$ (Fig. S21). As the recombination rate $r_{\mathrm{TC}}$ decreases, intermediate choosiness maximizing divergence at the T locus remains favored, but the magnitude of indirect selection favoring intermediate choosiness decreases, whether such intermediate choosiness evolves from low or high choosiness (Fig. S22a). This means that any other evolutionary force in the system favoring the evolution of high choosiness would be more likely to prevail as $r_{\mathrm{TC}}$ decreases. There is no such evolutionary force in the case of a magic trait (a change in $r_{\mathrm{TC}}$ affects the magnitude of the selection affecting choosiness, but not the direction of selection), unlike in the case of a nonmagic trait complex that we now examine.

With a nonmagic trait complex, the build-up of three-way linkage disequilibrium ultimately favors the evolution of high choosiness (as explained in the main text), but indirect viability and sexual selection also favor intermediate choosiness just like with a magic trait (as explained in the previous paragraph). Again, as the recombination rate $r_{\mathrm{TC}}$ decreases, the magnitude of indirect selection favoring intermediate choosiness decreases, whether such intermediate choosiness evolves from low or high choosiness. Therefore, as the recombination rate $r_{\mathrm{TC}}$ decreases, the constraint on the evolution of very high choosiness, higher than the choosiness that maximizes divergence, is reduced, and the effect of three-way linkage disequilibrium on the evolution of high choosiness can thus overcome this constraint (Fig. S22b).

Therefore, with a nonmagic trait complex, the evolutionary force favoring the evolution of very high choosiness (via the build-up of a three-way linkage disequilibrium) prevails mainly when the magnitude of indirect selection favoring intermediate choosiness (i.e., constraining the evolution of high choosiness) decreases. This occurs as the recombination rate $r_{\mathrm{TC}}$ decreases.

#### With weak costs of choosiness, pseudomagic trait complexes promote the evolution of higher choosiness than magic traits

Previous work has shown that costs to choosiness can be very significant in the context of sexual selection (e.g., they qualitatively change the outcomes of Fisherian runaway; Pomiankowski, 1987, *J. Theor. Biol.* **128**(2):195-218, and Bulmer, 1989, *Theor. Popul. Biol.* **35**(2):195-206). We thus consider here a variant of the model with selection on choosiness in the form of costs. The magnitude of these costs may or may not depend on the composition of the population; for example, mechanistic costs should remain the same (a "fixed" cost), but search costs should decline as the relative frequency of preferred mates increases (a "relative" cost). Following Otto et al. (2008, *Genetics* **179**(4):2091-2112), we include these two types of costs in our population genetics model, as detailed below (see also *Mathematica* notebook).

##### ***Preference matrix***

First, we consider a reparameterization of the preference matrix describing how females prefer to mate with certain males over others:

$$\begin{array}{ccc} & & \text{Male genotype} \\ & & \begin{array}{cc} T_1 & T_2 \end{array} \\ \begin{array}{c} \text{Female} \\ \text{genotype} \end{array} & \begin{array}{c} C_1T_1 \\ C_1T_2 \\ C_2T_1 \\ C_2T_2 \end{array} & \begin{pmatrix} 1 & 1-\rho_1 \\ 1-\rho_1 & 1 \\ 1 & 1-\rho_2 \\ 1-\rho_2 & 1 \end{pmatrix} \end{array} \tag{S17}$$

The terms $\rho_1$ and $\rho_2$ measure the degree to which a female dislikes males that do not match their T locus when she carries a $C_1$ or a $C_2$ allele, respectively. These terms $\rho_n$ depend on the choosiness parameters $\alpha_n$, so that

$$\rho_n = \frac{\alpha_n}{1+\alpha_n}. \tag{S18}$$

##### ***Fixed cost of choosiness***

We allowed for a fixed cost of choosiness, $c_{\rm f}$, which is paid by choosy females regardless of the types of males encountered.

After viability selection, the fitness of a female of genotype $i$ characterized by allele $C_{n_i}$ at the choosiness locus is multiplied by a fixed factor, so that

$$x_{i,k}^{**\rm f} = \frac{\left(1-\rho_{n_i}c_{\rm f}\right)x_{i,k}^{**}}{\sum_j \left(1-\rho_{n_j}c_{\rm f}\right)x_{j,k}^{**}}. \tag{S19}$$

Thus, at this point of the life cycle, the genotypic frequencies differ between males and females, and are represented by $x_{i,k}^{**}$ and $x_{i,k}^{**\rm f}$, respectively.

##### ***Relative cost of choosiness (search cost)***

We also allowed for a relative cost of assortment, $c_{\rm r}$, which is paid by a choosy female when she rejects a male.

Each female encounters a male and chooses to mate with him with a probability equal to the appropriate entry in matrix (S17). Equation S18 ensures that we get the same proportion of matings between females and males of particular genotypes as when we use Equation S2. In population $k$, the overall probability, summed over all possible types of males that a female with genotype $i$ characterized by alleles $T_{l_i}$ and $C_{n_i}$ accepts a male during a mating encounter, is

$$Q_{i,k} = t_{l_i,k}^{**} + \left(1-\rho_{n_i}\right)t_{m,k}^{**}, \tag{S20}$$

where $t^{**}_{l_i,k}$ and $t^{**}_{m,k}$ represent the frequency of males with allele $T_{l_i}$ and $T_m$ after viability selection occurred in population $k$. Here, $m = 2$ when $l_i = 1$, and $m = 1$ when $l_i = 2$.

If a female rejects a male, she may or may not be able to recuperate the lost mating opportunity. To account for this potential cost, we assume that a fraction of the time, $(1 - c_{\mathrm{r}})$, a female is able to recover the fitness lost by rejecting a dissimilar mate, and otherwise she suffers a loss in fitness. The overall chance that a female of genotype $i$ mates in population $k$ (which can be referred to as her "fertility") is then

$$F_{i,k} = (1 - c_{\mathrm{r}}) + c_{\mathrm{r}}\, Q_{i,k}. \tag{S21}$$

This cost of choosiness is relative; even a very picky female suffers no loss in fertility if she only encounters preferred males.

The average fertility of females in population $k$ is then

$$\bar{F}_k = \sum_i x^{**\mathrm{f}}_{i,k} \times F_{i,k}, \tag{S22}$$

with $x^{**\mathrm{f}}_{i,k}$ the frequency of females with genotype $i$ in population $k$ after choosy females paid a fixed cost (see previous subsection).

We then assume that, in population $k$, the frequency of mated pairs of females with genotype $i$ and males with genotype $j$ is not described by Equation S2, but by:

$$M_{ij,k} = \frac{F_{i,k}}{\bar{F}_k} \times \frac{(1 + \delta_{c_1,ij}\alpha_1)\,(1 + \delta_{c_2,ij}\alpha_2)\, x^{**}_{j,k}\, x^{**\mathrm{f}}_{i,k}}{\sum_z (1 + \delta_{c_1,iz}\alpha_1)\,(1 + \delta_{c_2,iz}\alpha_2)\, x^{**}_{z,k}}\,. \tag{S23}$$

The right term of this equation is the same as in Equation S2, at the exception of the consideration of the genotypic frequencies, $x^{**\mathrm{f}}_{i,k}$, in females after choosy females paid a fixed cost. The left term here accounts for the relative cost of choosiness controlled by parameter $c_{\mathrm{r}}$.

##### ***Effect of the genetic architecture on the ESS choosiness when choosiness is costly***

As in Figure 2, we determine numerically the ESS choosiness value in cases where choosiness evolves alongside a magic trait versus a nonmagic trait complex, but we here consider that female choosiness associates either with a fixed cost (controlled by $c_{\mathrm{f}} > 0$) or with a relative cost (controlled by $c_{\mathrm{r}} > 0$).

Fixed costs and relative costs have qualitatively the same effects on the ESS choosiness (top row vs. bottom row in Fig. S11). As expected, if the cost is strong enough, random mating is favored (e.g., for $r_{\mathrm{TC}} = 0.01$ or $= 0.05$ in the right column of Fig. S11; for higher costs, random mating is favored for $r_{\mathrm{TC}} = 0.5$ as well).

With a magic trait ($r_{\mathrm{ET}} = 0$), a low recombination rate between the choosiness locus and the pleiotropic locus (low $r_{\mathrm{TC}}$) leads to a very low ESS choosiness (Fig. S11). This is because a low $r_{\mathrm{TC}}$ reduces the constraint on the evolution of very high choosiness, as explained in the previous Supplementary Information Text. As the recombination rate $r_{\mathrm{TC}}$ decreases, the magnitude of indirect selection favoring intermediate choosiness decreases, and any evolutionary force inhibiting the evolution of choosiness, such as that caused by a cost of choosiness, is therefore more likely to prevail.

With a nonmagic trait ($r_{\mathrm{ET}} > 0$), the establishment of three-way linkage disequilibrium can favor the evolution of higher choosiness than the ESS choosiness established with a magic trait, even if weak costs of choosiness inhibit the evolution of choosiness (e.g., middle column in Fig. S11). Consequently, in some cases, the difference in ESS choosiness evolving alongside nonmagic and magic traits can even be amplified by a weak cost of choosiness (e.g., see differences in ESS choosiness when $r_{\mathrm{ET}} = 0$ vs. $r_{\mathrm{ET}} = 0.2$ in red, in Fig. S11). Note however that this does not occur in a diploid version of the model with similar costs (see next Supplementary Information Text).

Therefore, even with weak costs of choosiness, pseudomagic trait complexes can promote the evolution of stronger assortative mate choice than do magic traits, provided that polymorphism at the ecological and mating signal loci is maintained.

#### With a diploid life cycle, pseudomagic trait complexes can promote the evolution of higher choosiness than magic traits

We also develop a three-locus diploid model that is analogous to the haploid model presented in the main text. We invite the reader to consult the *Mathematica* notebook for details on all recursion equations. Below we summarize the changes we made due to the consideration of diploidy.

##### *Genotypes*

We consider three loci, E, T and C, that encode the same traits as in our haploid model. In our diploid model, however, there are $4^3 = 64$ genotypes. Just like in the haploid model, the life cycle is as follows: census, migration, viability selection, mating, production of zygotes.

##### *Migration*

Each generation first undergoes symmetric migration with rate $m$. We model the migration step as in our haploid model.

##### *Viability selection*

Next, divergent selection occurs. The ecological trait encoded by locus E is locally adapted, such that each ecological allele, $E_1$ and $E_2$, is favored by viability selection in the population in which it was common in allopatry. Due to the diploidy, we have to make additional assumptions. We assume that in population $k$, homozygotes $E_kE_k$ have a selective advantage due to local adaptation, with relative fitness $1 + s_k$. For simplicity, we also consider that both heterozygotes $E_1E_2$ and $E_2E_1$ have a relative fitness $1 + \beta\, s_k$, where the parameter $\beta \in [0, 1]$ measures the degree of deme-dependent dominance (such that in deme $k$, $E_k$ is dominant for $\beta = 1$, and recessive for $\beta = 0$). Parameter $\beta$ thus reflects the extent to which heterozygotes at the E locus are on average more or less fit than homozygotes. We assume symmetric viability selection such that $s_1 = s_2 = s$.

##### *Mating*

Mating follows, during which females express different propensities to mate with males. In our diploid model, we again have to impose additional assumptions over the haploid version. Specifically, heterozygous females $C_nC_m$ that are also homozygous $T_1T_1$ (resp. $T_2T_2$) are $1+\alpha_{nm}$ times as likely to mate with a $T_1T_1$ male (resp. a $T_2T_2$ male) than with a male carrying only the opposite $T$ alleles, if she encounters one of each. Additionally, these females are $1+\gamma\,\alpha_{nm}$ times as likely to mate with an heterozygous male $T_1T_2$ or $T_2T_1$. The parameter $\gamma \in [0,1]$ thus measures the inability of a female to distinguish males that differ by one vs. two alleles at the T locus, when she prefers her own genotype.

We also assume that $\alpha_{12} = \alpha_{21} = \alpha_{11} + h_{\mathrm{C}}\,(\alpha_{22} - \alpha_{11})$ with the parameter $h_{\mathrm{C}} \in [0,1]$ being the dominance coefficient at the C locus.

In the results shown in the subsequent figures, we assume that heterozygous females $T_1T_2$ and $T_2T_1$ have no preference. In other sets of simulations shown in the *Mathematica* notebook, however, we assumed that heterozygous females $T_1T_2$ and $T_2T_1$ express mating preferences. Specifically, heterozygous females $C_nC_m$ that are also heterozygous at the T locus are $1+\alpha_{nm}$ times as likely to mate with another heterozygote at the T locus. Additionally, these females are $1+\gamma\,\alpha_{nm}$ times as likely to mate with an homozygous male $T_1T_1$ or $T_2T_2$. If heterozygotes at the T locus are frequent in the population, this can generate stabilizing sexual selection on the T locus. Given that this does not change qualitatively our results, we do not show the graphs here (but see *Mathematica* notebook).

We also consider selection on choosiness in the form of fixed and relative costs, controlled by parameters $c_{\mathrm{f}}$ and $c_{\mathrm{r}}$, as detailed in the previous Supplementary Information Text. The frequency of mated pairs of females and males with particular genotypes is then described by Equation S23.

##### *Production of zygotes*

Males and females produce haploid gametes during meiosis. During meiosis, recombination occurs at a rate $r_{\mathrm{ET}}$ between loci E and T, and $r_{\mathrm{TC}}$ between loci T and C. In each population, the expected genotype frequencies of diploid zygotes in the next generation are therefore calculated by summing the appropriate mating frequencies assuming Mendelian segregation and recombination during the production of haploid gametes.

##### *Effect of the genetic architecture on the ESS choosiness in the diploid model*

With the diploid version of our model, we determine numerically the ESS choosiness value in cases where choosiness evolves alongside a magic trait versus a pseudomagic trait complex, as was done for the haploid model in Figure 2.

We obtain qualitatively the same outcome as in the haploid case: nonmagic trait complexes can promote the evolution of stronger assortative mate choice than do magic traits, provided that polymorphism at the ecological and mating signal loci is maintained (Fig. S12). Note, however, that selection against heterozygous hybrids strongly affects the ESS choosiness in our diploid version of the model. In particular, weak viability selection against hybrids (high $\beta$) decreases the ESS choosiness and reduces the effect of pseudomagic trait architectures. The former effect is driven by lower indirect selection favoring choosiness, and the latter effect is explained by a smaller contribution of the three-way linkage disequilibrium on the evolution of choosiness. Similar effects result from the inability of females to distinguish males that differ by one vs. two alleles at the T locus ($\gamma = 0.7$ in Fig. S12; the ESS choosiness actually reaches 0 for $\gamma = 1$). Note that for low $\gamma < 0.5$, high choosiness is consistently favored (see *Mathematica* notebook).

Unlike the parameters $\beta$ and $\gamma$, the dominance coefficient $h_{\mathrm{C}}$ at the C locus has no effect at all on the ESS choosiness (although the speed of invasion of recessive choosiness alleles that are beneficial is reduced compared to dominant ones; not shown, but see *Mathematica* notebook).

Interestingly, the effects of costs of choosiness we highlighted in the previous Supplementary Information Text do not occur in the diploid version of the model. The difference in ESS choosiness evolving alongside nonmagic and magic traits is not amplified by a weak cost of choosiness (Fig. S13). This is because in the haploid model, selection favoring intermediate choosiness is very weak for low $r_{\mathrm{TC}}$, whereas in the diploid model, selection against maladapted hybrids occurs even for low $r_{\mathrm{TC}}$.

To sum up, even with a diploid life cycle, pseudomagic trait complexes can promote the evolution of stronger assortative mate choice than do magic traits.

### Supplementary Figures

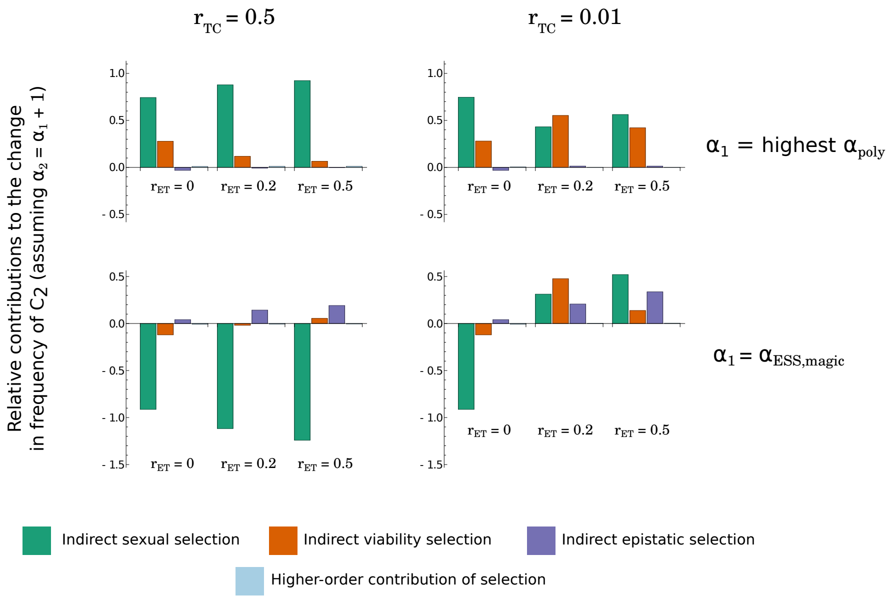


**Figure S1: Relative first-order contributions to the evolution of stronger choosiness of sexual selection, viability selection, epistatic selection, and higher-order contribution of selection.** We consider the invasion of an allele coding for higher choosiness, $C_2$, such that $\alpha_2 = \alpha_1 + 1$, and represent the mean first-order contributions of linkage disequilibria to the change in frequency of the $C_2$ allele in population 2, $\Delta c_{2,2}$, for different recombination rates ($r_{\mathrm{ET}}$ and $r_{\mathrm{TC}}$) and different choosiness values ($\alpha_1$). In the top row, choosiness $\alpha_1$ is set to be the lowest choosiness value that maintains polymorphism at the T locus for all recombination rates tested ($\alpha_1$ = 'highest $\alpha_{\mathrm{poly}}$' $= 0.13$; estimated numerically). In the bottom row, choosiness $\alpha_1$ is set to be the ESS choosiness value obtained for $r_{\mathrm{ET}} = 0$ ($\alpha_1 = \alpha_{\mathrm{ESS,magic}} = 6.54$; see Figure 2). For each combination of parameters $(r_{\mathrm{ET}}, r_{\mathrm{TC}}, \alpha_1)$, we know if a choosier allele $C_2$ will increase or decrease in frequency based on the ESS choosiness value. We therefore implement $C_2$ at a frequency equal to 0.01 if it is destined to increase in frequency, or equal to 0.99 if it is destined to decrease in frequency. Over the course of the simulation, while the frequency of the choosier allele $c_{2,2}$ is between 0.05 and 0.95, we measure the mean first-order contributions of linkage disequilibrium to $\Delta c_{2,2}$, corresponding to the first three terms of Equation 1, and the mean higher-order contribution of linkage disequilibrium (in light blue), corresponding to the last term in Equation 1. These first order-contributions of linkage disequilibria correspond to first-order approximations of the effect of indirect sexual selection (in green), indirect viability selection (in orange), and indirect epistatic selection (in purple) on the change in frequency of the choosier allele. We then normalize these contributions to measure their relative contribution to the change in frequency of allele $C_2$. Note that we can draw the same conclusions by tracking changes in frequencies in population 1 (not shown). In the case of a magic trait (for $r_{\mathrm{ET}} = 0$), the first three contributions can be summed together to assess the first-order approximation of indirect selection acting on the choosiness modifier via linkage disequilibrium with the pleiotropic locus, which fulfills the functions of both E and T loci. For each combination of parameters, the sum of the four contributions equals to one. See more explanations on the effect of $r_{\mathrm{ET}}$ on these contributions in Fig. S2. Here, $m = 0.01$ and $s = 0.05$.

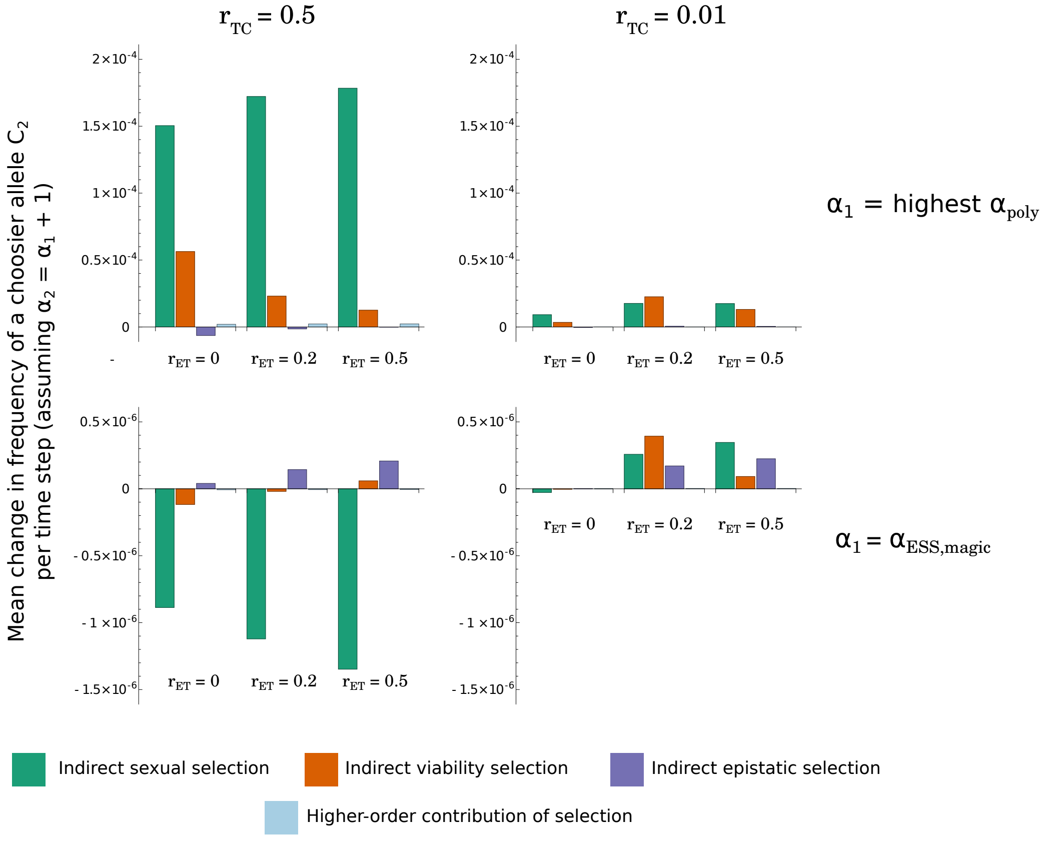


**Figure S2: First-order contributions to the evolution of stronger choosiness of sexual selection, viability selection, epistatic selection, and higher-order contribution of selection.** We consider the same simulations as in Fig. S1 (see caption for more details), but this time we represent the actual mean contributions of these selective forces to the evolution of stronger choosiness (we do not apply normalization). Below we describe how $r_{\mathrm{ET}}$ affect these contributions, so that the reader can better understand Fig. 3b.

A high $r_{\mathrm{ET}}$ slightly inhibits divergence at the T locus (Fig. 1b); this leads to an increase in linkage disequilibrium $D_{\mathrm{TC}}$ because there is more variation at the underlying loci, thereby **increasing the contribution of sexual selection** to the evolution of choosiness (green bars; although this slight decrease in divergence at the T locus reduces the strength of frequency-dependent sexual selection).

A high $r_{\mathrm{ET}}$ strongly inhibits divergence at the E locus (Fig. 1b); this leads to an increase in linkage disequilibrium $D_{\mathrm{EC}}$. Nonetheless, recombination occurring at a rate $r_{\mathrm{ET}}$ also directly degrades linkage disequilibrium, $D_{\mathrm{EC}}$. This last effect prevails explaining why a high $r_{\mathrm{ET}}$ **decreases the contribution of viability selection** to the evolution of choosiness (orange bars; in particular for $r_{\mathrm{TC}} = 0.5$). This effect can be offset when the build-up of three-way linkage disequilibrium, allowed by recombination between the E and T loci, leads to the build-up of $D_{\mathrm{EC}}$ (for $r_{\mathrm{TC}} = 0.01$). In that case a high $r_{\mathrm{ET}}$ **increases the contribution of viability selection** to the evolution of choosiness. Finally, as detailed in the main text, a high $r_{\mathrm{ET}}$ contributes to the build-up of three-way linkage disequilibrium, thereby **increasing the contribution of epistatic selection** to the evolution of choosiness (purple bars).

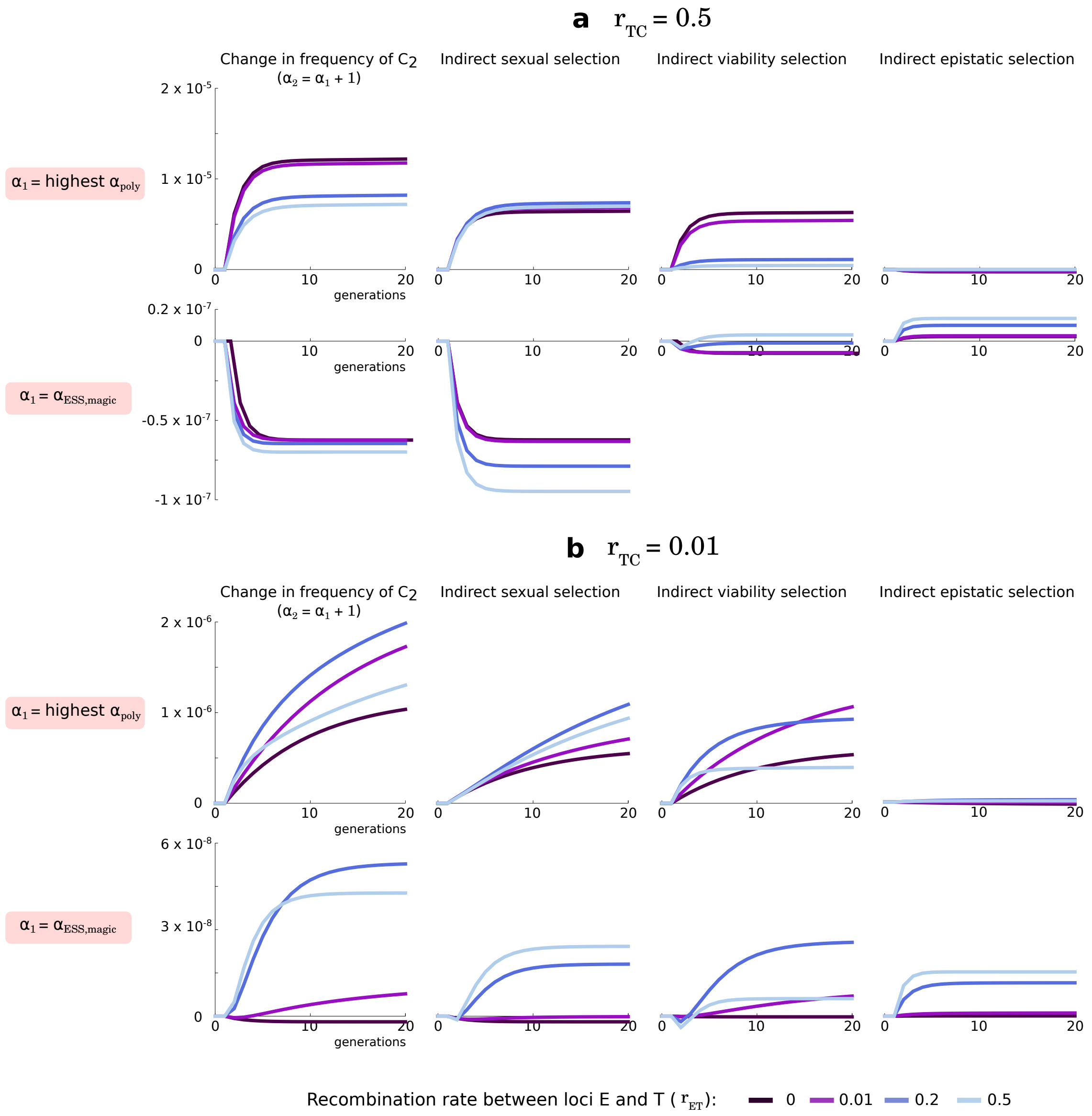


**Figure S3: Effect of indirect selection over time on the change in frequency of a choosier allele.** We represent the change in frequency per time step, $\Delta c_{2,k}$, of a choosier allele, and the first order contributions of indirect sexual selection, indirect viability selection, and indirect epistatic selection (the first three terms of Equation 1), to the evolution of stronger choosiness for different recombination rates ($r_{\mathrm{ET}}$ and $r_{\mathrm{TC}}$) and different choosiness values ($\alpha_1$ and $\alpha_2$). In the top rows of panels **a** and **b**, choosiness $\alpha_1$ is set to be the choosiness value that maintains polymorphism at the T locus for all recombination rates tested ($\alpha_1$ = 'highest $\alpha_{\mathrm{poly}}$' = 0.13; estimated numerically). In the bottom rows of panels **a** and **b**, choosiness $\alpha_1$ is set to be the ESS choosiness value obtained for $r_{\mathrm{ET}} = 0$ ($\alpha_1 = \alpha_{\mathrm{ESS,magic}} = 6.54$; see Figure 2). Because the signs of the linkage disequilibria can be inferred from the directions of the contributions of indirect selection, the time series of linkage disequilibria are qualitatively similar to these time series (Fig. S4). Note that in the bottom row of panel **b**, indirect sexual and viability selection eventually favor the evolution of higher choosiness, although they inhibit the evolution of higher choosiness during the first couple of generations. This is because during the first couple of generations, three-way linkage disequilibrium has not yet generated two-way linkage disequilibria, leading to positive indirect sexual and viability selection. Here, $m = 0.01$ and $s = 0.05$.

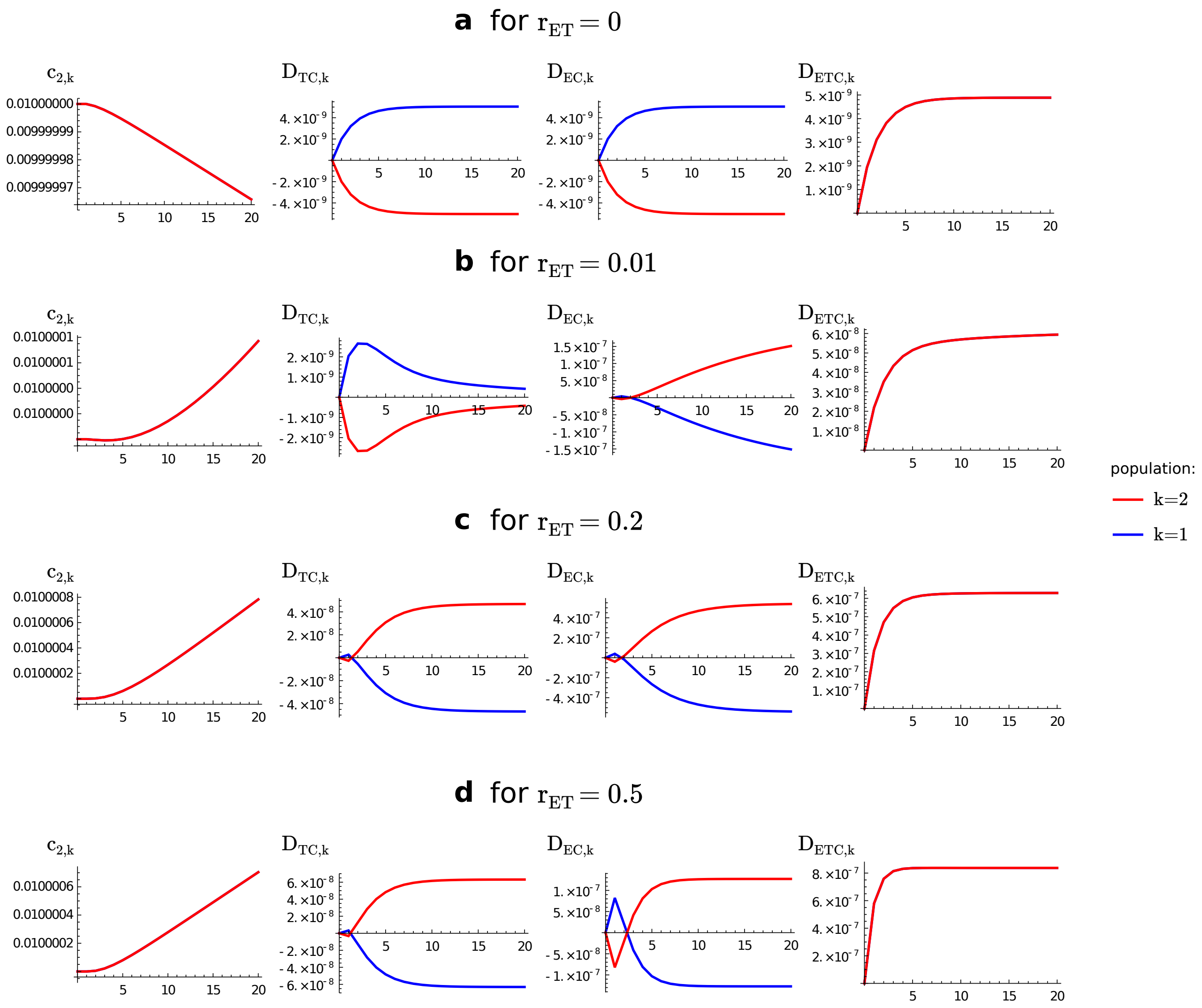


**Figure S4: Time series of allelic frequency at the choosiness locus and linkage disequilibria.** We represent the allelic frequency at the choosiness locus ($c_{2,k}$), pairwise linkage disequilibria involving the choosiness locus ($D_{\mathrm{TC},k}$, $D_{\mathrm{EC},k}$), and three-way linkage disequilibrium ($D_{\mathrm{ETC},k}$) in each population $k$, for different recombination rate $r_{\mathrm{ET}}$. Here, we consider the same conditions as in the bottom row of Figure S3 (i.e., for $r_{\mathrm{TC}} = 0.01$, and $\alpha_1$ set to be the ESS choosiness value obtained for $r_{\mathrm{ET}} = 0$; see caption of Fig. S3 for details). In the far left and far right columns, the red and blue lines overlap. As already shown in Figure S3, we observe here that pairwise linkage disequilibria favoring a choosier allele $C_2$ can take a couple of generations to become established (because it takes a bit of time for three-way linkage disequilibrium to be high enough so that it can lead to the establishment of pairwise linkage disequilibria that ultimately favor $C_2$). This dynamics of linkage disequilibria occurs for any nonmagic trait ($r_{\mathrm{ET}} > 0$; **b-d**), but it is particularly visible for $r_{\mathrm{ET}} = 0.5$ (**d**).

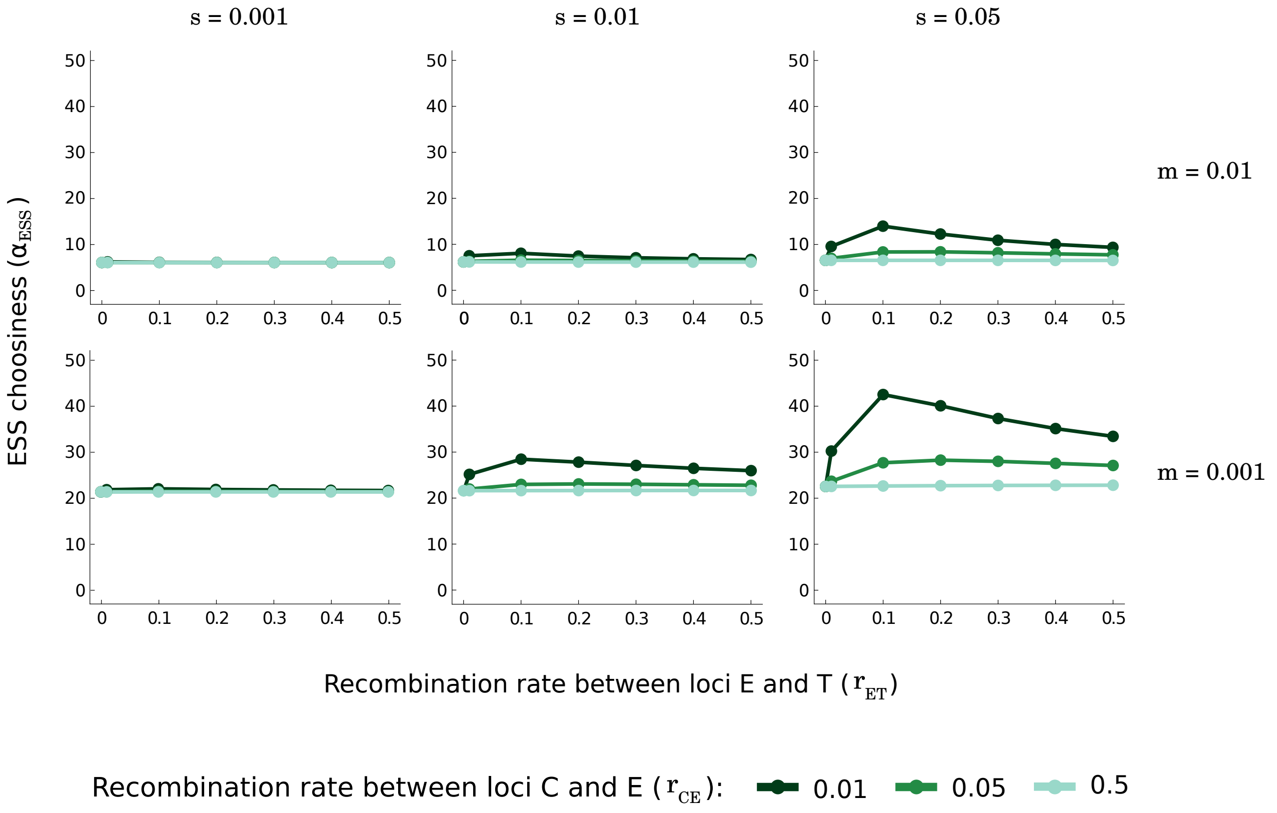


**Figure S5: Choosiness at evolutionary equilibrium depending on the level of gene flow and genetic architecture when the gene order is CET.** We implement the same combinations of parameters as in Fig. 2, but we vary the recombination rate $r_{\mathrm{CE}}$ between loci C and E, instead of the recombination rate between loci T and C. With gene order CET, we necessarily have $r_{\mathrm{ET}} < r_{\mathrm{TC}}$, and therefore the conditions where the ESS choosiness is the highest with gene order ETC ($r_{\mathrm{TC}} = 0.01$ and $r_{\mathrm{ET}} > r_{\mathrm{TC}}$, Fig. 2) cannot be met with gene order CET. That is why the ESS choosiness obtained with the gene order CET is lower than that obtained with gene order ETC. Overall, however, we get qualitatively the same outcome. Just like with gene order ETC, for a given combination of parameters $(s, m)$, an intermediate $r_{\mathrm{ET}}$ leads to the highest ESS choosiness. For $r_{\mathrm{CE}} = 0.5$, changes in the recombination rate $r_{\mathrm{ET}}$ lead to slight changes in the ESS choosiness that are not visible here, just like with gene order ETC. We do not consider gene order ECT, because in that case, the choosiness locus would necessarily be fully linked to the other loci if we consider a magic trait (when $r_{\mathrm{ET}} = 0$). In the case of pseudomagic trait complexes, however, we can expect that gene order ECT would lead to high ESS choosiness, because the choosiness locus would necessarily be linked to the other loci (because $r_{\mathrm{TC}}$ and $r_{\mathrm{EC}}$ need to be smaller than $r_{\mathrm{ET}}$), as explained in the main text.

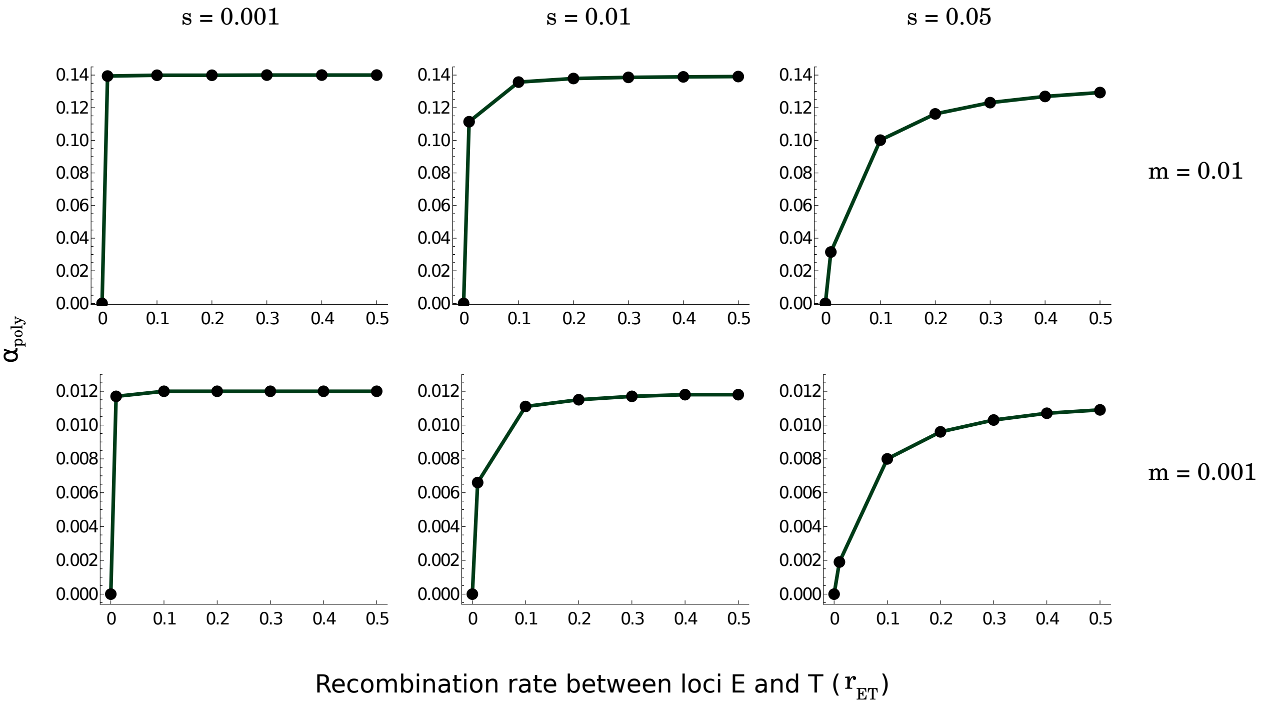


**Figure S6: Lowest choosiness value, $\alpha_{\mathbf{poly}}$, allowing the maintenance of polymorphism at evolutionary equilibrium depending on the level of gene flow and genetic architecture.** We implement the same combinations of parameters as in Fig. 2 ($r_{\mathrm{TC}}$ has no effect because we assume here that there is no variation in choosiness), and we measure $\alpha_{\mathrm{poly}}$. Magic traits allow the maintenance of polymorphism under random mating. In contrast, nonmagic trait complexes allow the maintenance of polymorphism only under nonrandom mating ($\alpha_{\mathrm{poly}} > 0$). Note that $\alpha_{\mathrm{poly}}$ remains very small. Therefore, polymorphism is maintained alongside nonmagic trait complexes if at least a small level of choosiness has already evolved before secondary contact occured.

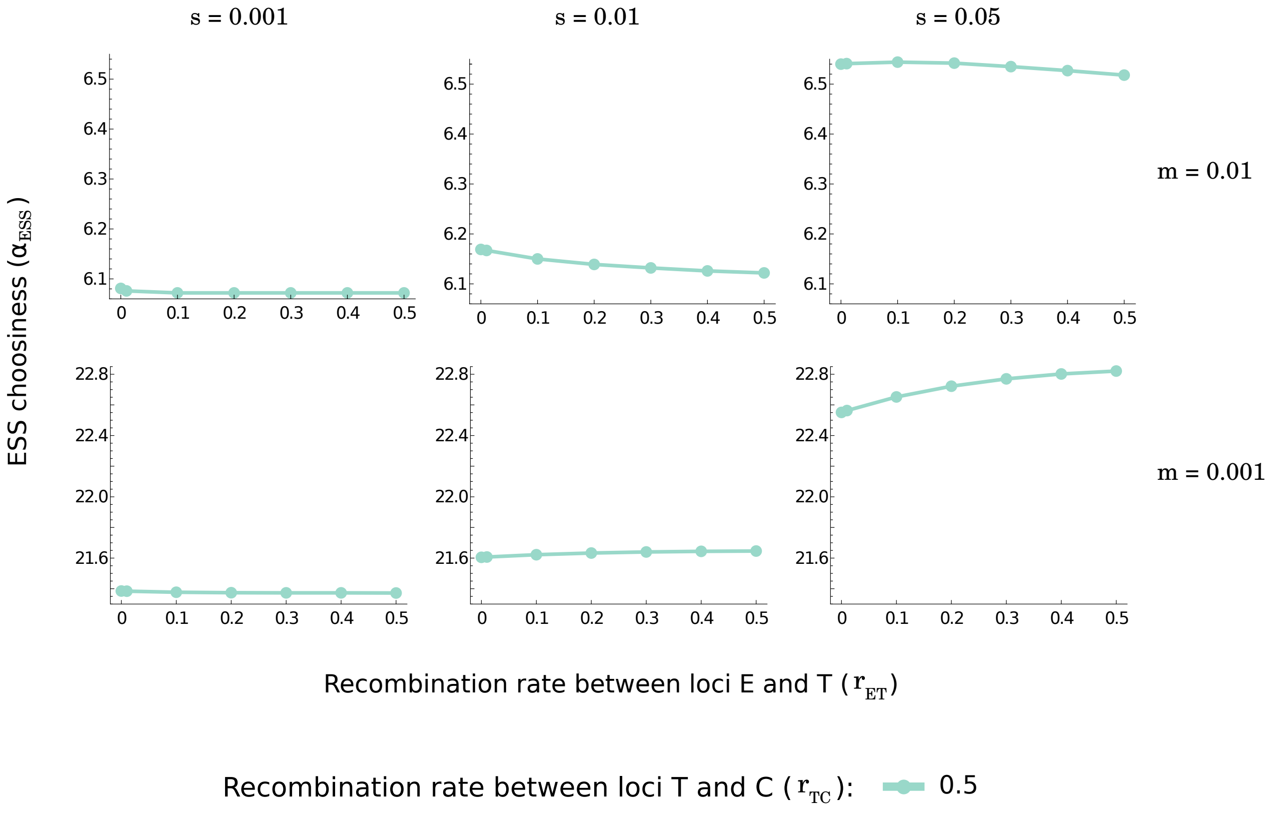


**Figure S7: Choosiness at evolutionary equilibrium depending on the level of gene flow and genetic architecture when $\mathbf{r_{TC} = 0.5}$.** We implement the same combinations of parameters $(s, m)$ as in Fig. 2, but use different vertical axes to highlight how changes in the recombination rate $r_{\mathrm{ET}}$ lead to slight changes in ESS choosiness, $\alpha_{\mathrm{ESS}}$, when $r_{\mathrm{TC}} = 0.5$. In particular, for a high ratio $s/m$ (e.g., for $s = 0.05$ and $m = 0.001$), a nonmagic trait complex ($r_{\mathrm{ET}} > 0$) can lead to a higher choosiness at evolutionary equilibrium than can be found with a magic trait ($r_{\mathrm{ET}} = 0$). Those are the conditions that favor the evolution of high choosiness alongside divergence at a nonmagic trait complex (Fig. 2). A high $r_{\mathrm{ET}}$ may lead, under some conditions, to a lower ESS choosiness; this is caused by a lower contribution of indirect viability selection to the evolution of high choosiness, due to the fact that a high $r_{\mathrm{ET}}$ degrades linkage disequilibrium $D_{\mathrm{EC}}$ (see details in Figs. S1 and S2).

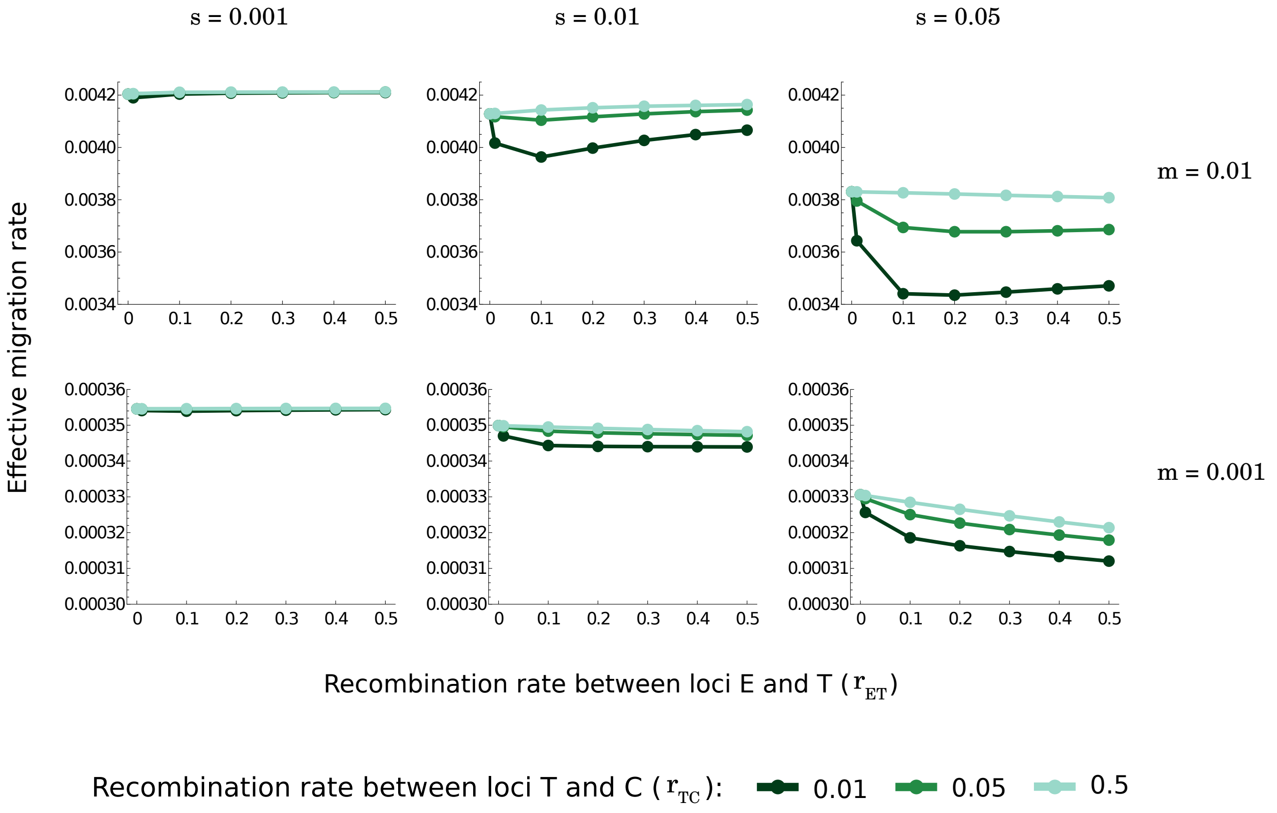


**Figure S8: Effective migration rate at the ESS choosiness depending on the level of gene flow and genetic architecture.** We implement the same combinations of parameters as in Fig. 2, and we numerically measure the effective migration rate at the ESS choosiness (a measure of gene flow) for a neutral locus that is linked to the trait loci. We adapt the approach of Akerman and Bürger (2014, *Theoretical Population Biology* **94**(100):42-62), who derived an eigenvalue-effective migration rate in a two-deme model for a neutral site linked to one or two loci. See Akerman and Bürger (2014), and Servedio and Bürger (2020, *Evolution* **74**(11):2438-2450) for details on the method. We here consider a neutral locus that is linked to the T locus, so that recombination occurs at a rate equal to 0.1 between the neutral locus and the T locus. In terms of interpreting the effective migration rate, one should compare its value to the migration rate to infer the effect of the ESS choosiness on reproductive isolation. A nonmagic trait complex leads to a higher ESS choosiness than a magic trait (Fig 2), but also leads to lower divergence at the T locus (e.g., Fig. 1; see also Servedio and Bürger, 2020). This lower divergence at the T locus increases the proportion of encounters between $T_1$ and $T_2$ individuals, which has the effect of reducing premating isolation. Overall, however, we show here that the increase in the ESS choosiness alongside a nonmagic trait complex prevails, out of these two effects, and leads to a net increase in premating isolation (i.e., it decreases the effective migration rate over a magic trait). Note that the vertical axis does not start at 0, and that the range on the vertical axes are different in the top and bottom rows. For even weaker selection relative to migration, there is loss of divergence, but those conditions are not shown.

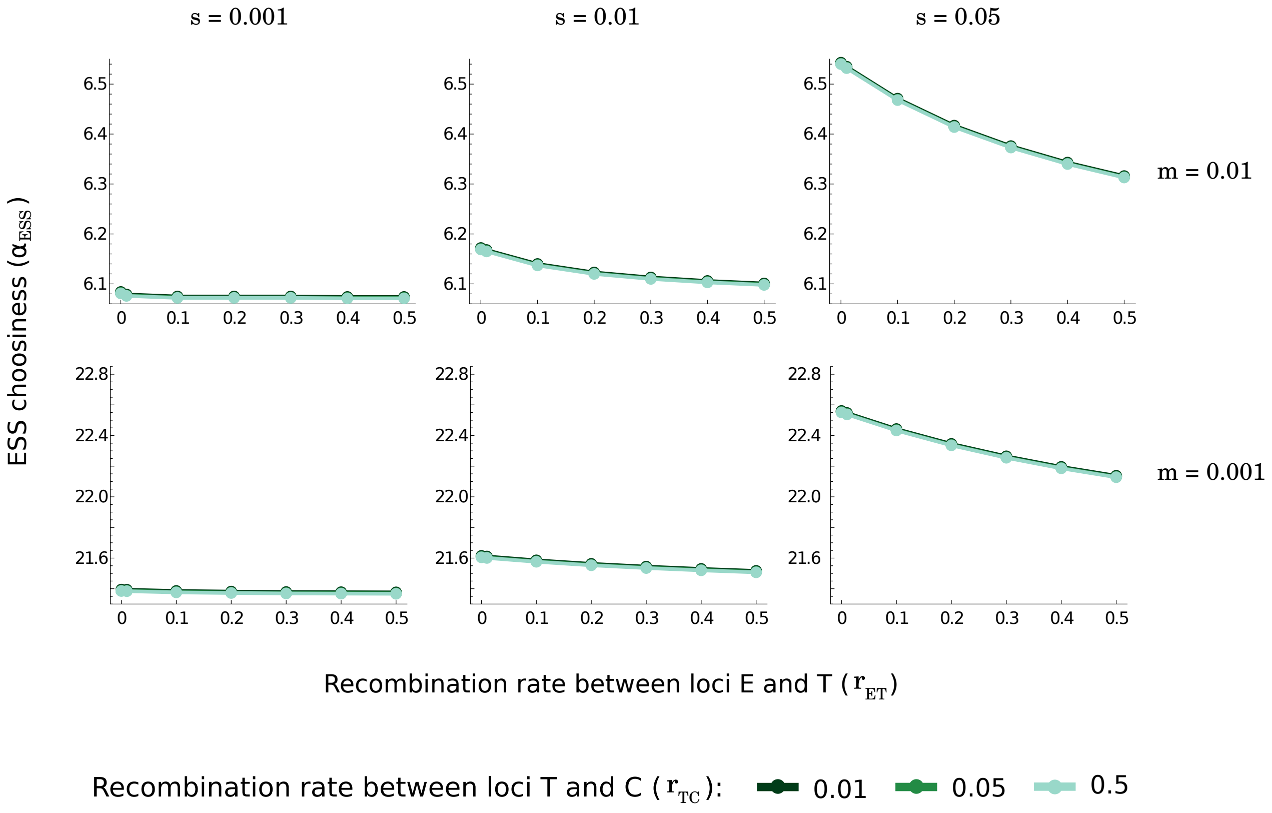


**Figure S9: Choosiness at evolutionary equilibrium depending on the level of gene flow and genetic architecture when the magnitude of the three-way linkage disequilibrium is artificially reduced to its lowest possible value.** We implement the same combinations of parameters as in Fig. 2. Over the course of simulations, however, here we artificially reduce the magnitude of the three-way linkage disequilibrium at the end of each generation ($\Phi = 1$; see caption of Fig. 3 for more details). In each panel, all lines and points overlap. This means that the ESS choosiness does not depend on the recombination rate $r_{\text{TC}}$ if we prevent the three-way linkage disequilibrium from building. We also note that if we prevent the three-way linkage disequilibrium from building, a nonmagic trait complex ($r_{\text{ET}} > 0$) consistently leads to a lower choosiness at evolutionary equilibrium than that of a magic trait ($r_{\text{ET}} = 0$).

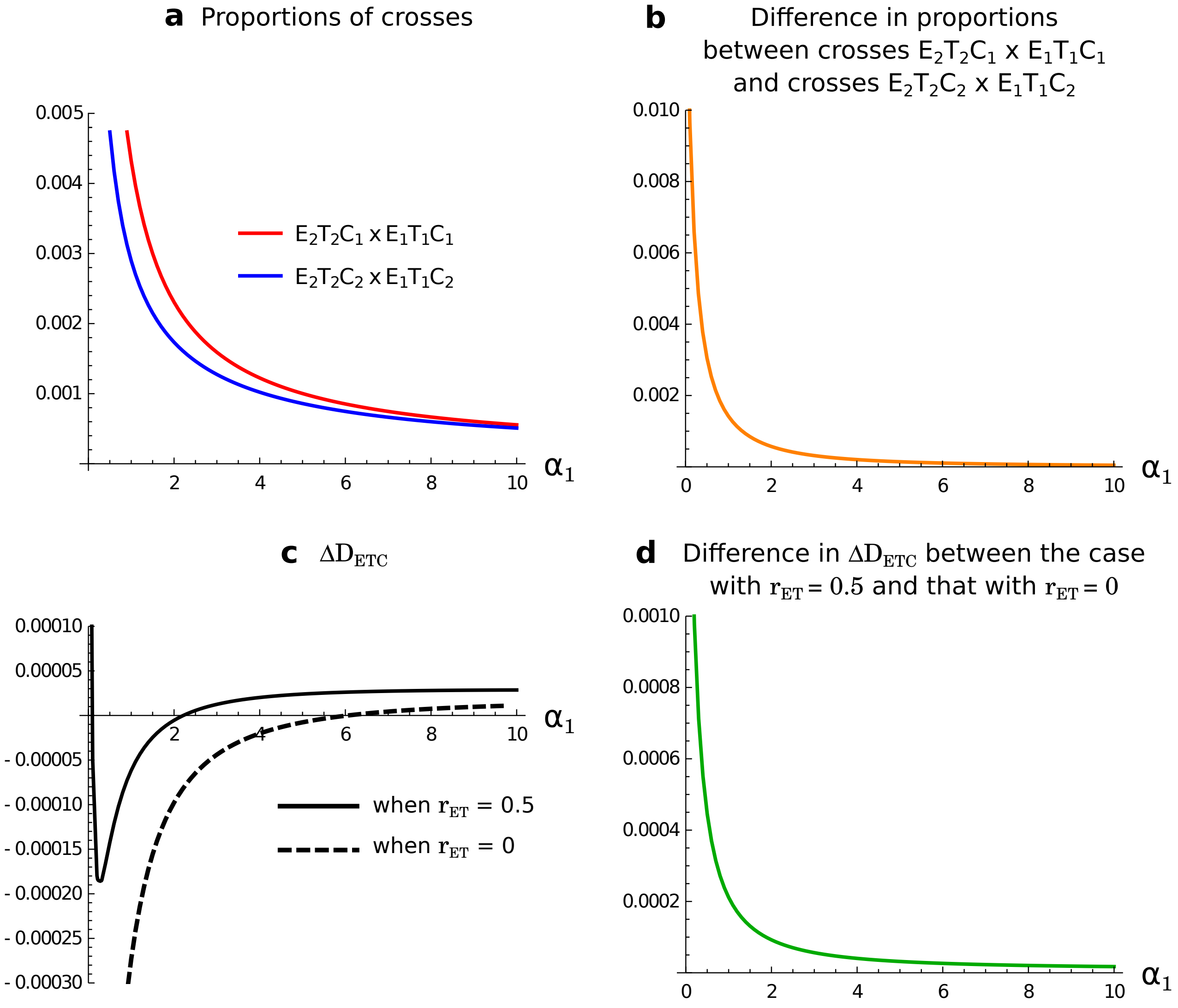


**Figure S10: Effect of the recombination rate between loci E and T on the build-up of three-way linkage disequilibrium among loci E, T and C.** We consider that allele $C_1$ codes for choosiness $\alpha_1$, and that allele $C_2$ codes for higher choosiness, $\alpha_2 = \alpha_1 + 1$. We start the population at the migration-selection equilibrium at the E and T loci given the ancestral choosiness, which is encoded by allele $C_1$. We introduce the allele $C_2$, coding for a higher choosiness, in linkage equilibrium with the other loci and in the same frequency ($= 0.01$) in both populations, and we allow migration and viability selection to occur. We then show, in population 2, (**a**) the proportions of crosses $E_2T_2C_1 \times E_1T_1C_1$ (red) and $E_2T_2C_2 \times E_1T_1C_2$ (blue) that break beneficial association $E_2T_2$ associated with alleles $C_1$ and $C_2$ respectively, via recombination between loci E and T, (**b**) the difference in proportions between these crosses, (**c**) the change in three-way linkage disequilibrium, $\Delta D_{\mathrm{ETC}}$, during mating and the production of zygotes with or without recombination between loci E and T (where the equivalent measure of $D_{\mathrm{ETC}}$ in the case of a magic trait is measured as $D_{\mathrm{TC,k}}(1 - 2t_{2,k})$ with T being the pleiotropic locus; see *Mathematica* notebook), and (**d**) the difference between these changes in three-way linkage disequilibrium $\Delta D_{\mathrm{ETC}}$ and its equivalent in the case of a magic trait. Here, $m = 0.01$, $s = 0.05$, and $r_{\mathrm{TC}} = 0.5$.

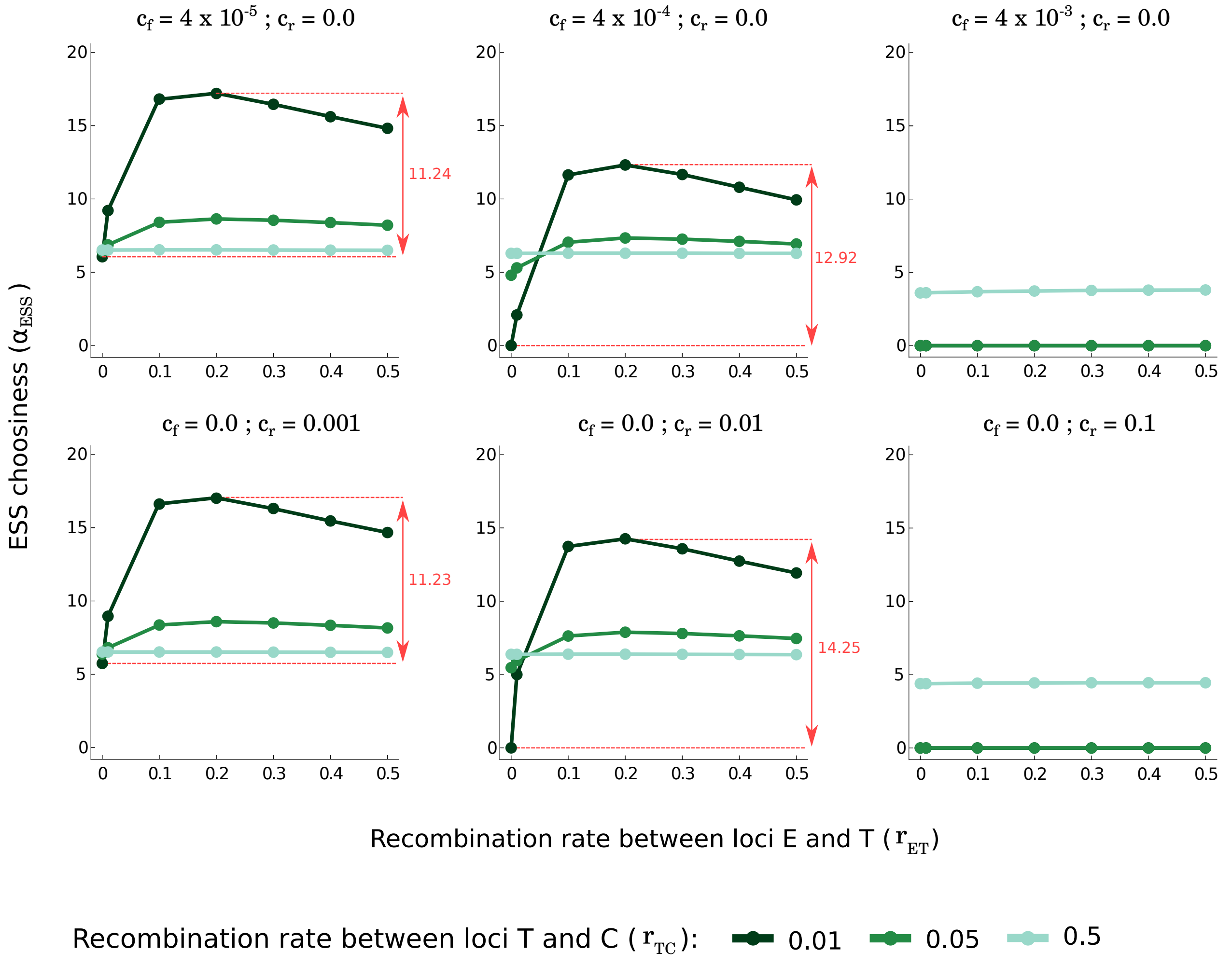


**Figure S11: Choosiness at evolutionary equilibrium depending on the level of gene flow and genetic architecture when choosiness is costly.** We represent the evolutionary stable choosiness, $\alpha_{\text{ESS}}$, depending on the recombination rates ($r_{\text{ET}}$ and $r_{\text{TC}}$), and the cost of choosiness (fixed and relative costs, controlled by $c_{\text{f}}$ and $c_{\text{r}}$, respectively). In red, we show the difference in ESS choosiness when $r_{\text{ET}} = 0$ vs. $r_{\text{ET}} = 0.2$, for $r_{\text{TC}} = 0.01$. Without cost ($c_{\text{f}} = 0$ and $c_{\text{r}} = 0$), this difference in ESS choosiness is equal to 11.21. See caption of Fig. 2 for more details. Here, $m = 0.01$ and $s = 0.05$.

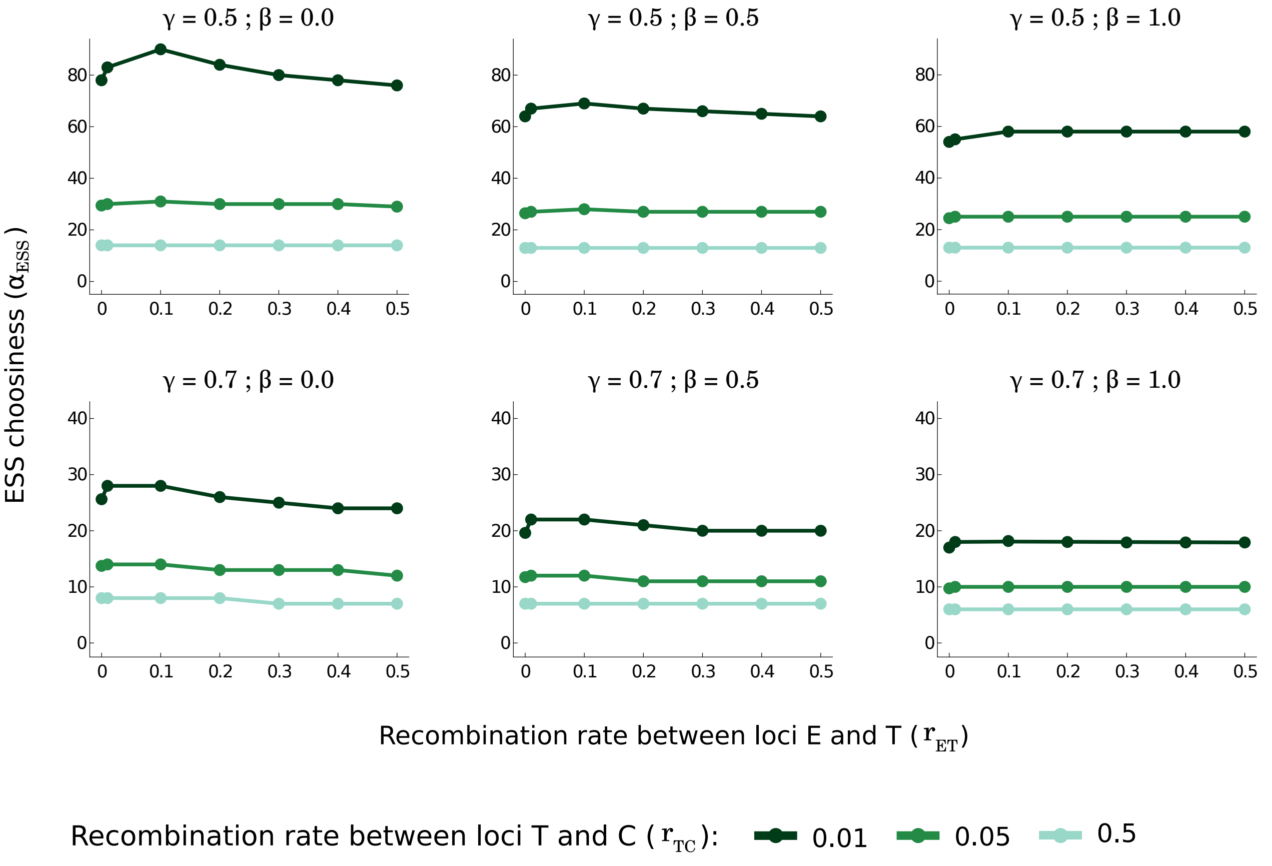


**Figure S12: Choosiness at evolutionary equilibrium depending on the level of gene flow and genetic architecture in the diploid version of the model.** We represent the evolutionary stable choosiness, $\alpha_{\text{ESS}}$, depending on the recombination rates ($r_{\text{ET}}$ and $r_{\text{TC}}$), the extent to which heterozygotes at the E locus are fit (controlled by $\beta$), and the inability of females to distinguish males that differ by one vs. two alleles at the T locus (controlled by $\gamma$). Here, we assume that heterozygotes at the T locus do not express mating preferences. When we assume that heterozygous females $T_1T_2$ and $T_2T_1$ express mating preferences, the ESS choosiness is slightly lower, but we get qualitatively the same outcome (not shown here; but see *Mathematica* notebook). We also do not include costs of choosiness ($c_{\text{f}} = 0$ and $c_{\text{r}} = 0$). See caption of Fig. 2 for more details. Here, $m = 0.01$ and $s = 0.05$.

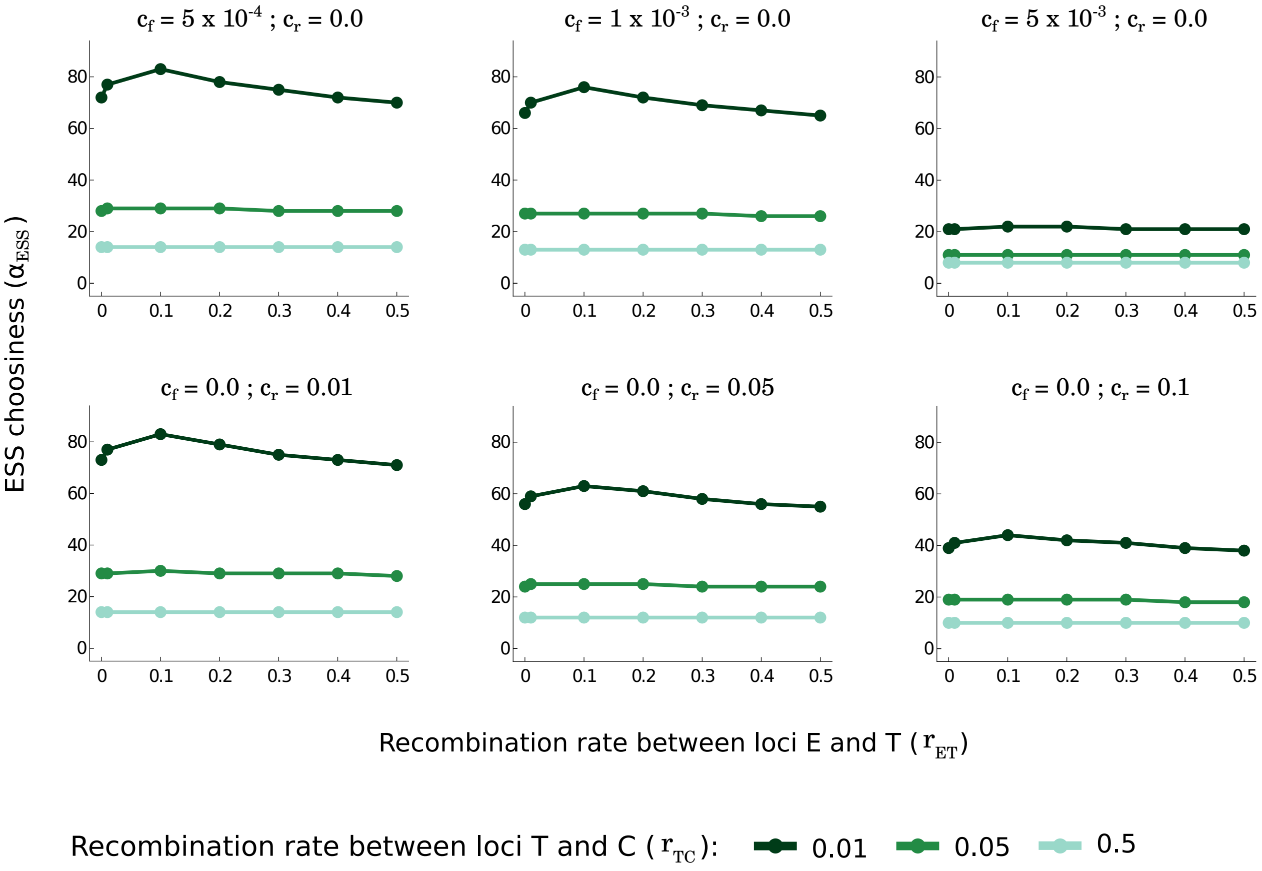


**Figure S13: Choosiness at evolutionary equilibrium depending on the level of gene flow and genetic architecture when choosiness is costly in the diploid version of the model.** We represent the evolutionary stable choosiness, $\alpha_{\text{ESS}}$, depending on the recombination rates ($r_{\text{ET}}$ and $r_{\text{TC}}$), and the cost of choosiness (fixed and relative costs, controlled by $c_{\text{f}}$ and $c_{\text{r}}$, respectively). Here, we assume that $\beta = 0$, $\gamma = 0.5$ and that heterozygotes at the T locus do not express mating preferences. See caption of Fig. 2 for more details. Here, $m = 0.01$ and $s = 0.05$.

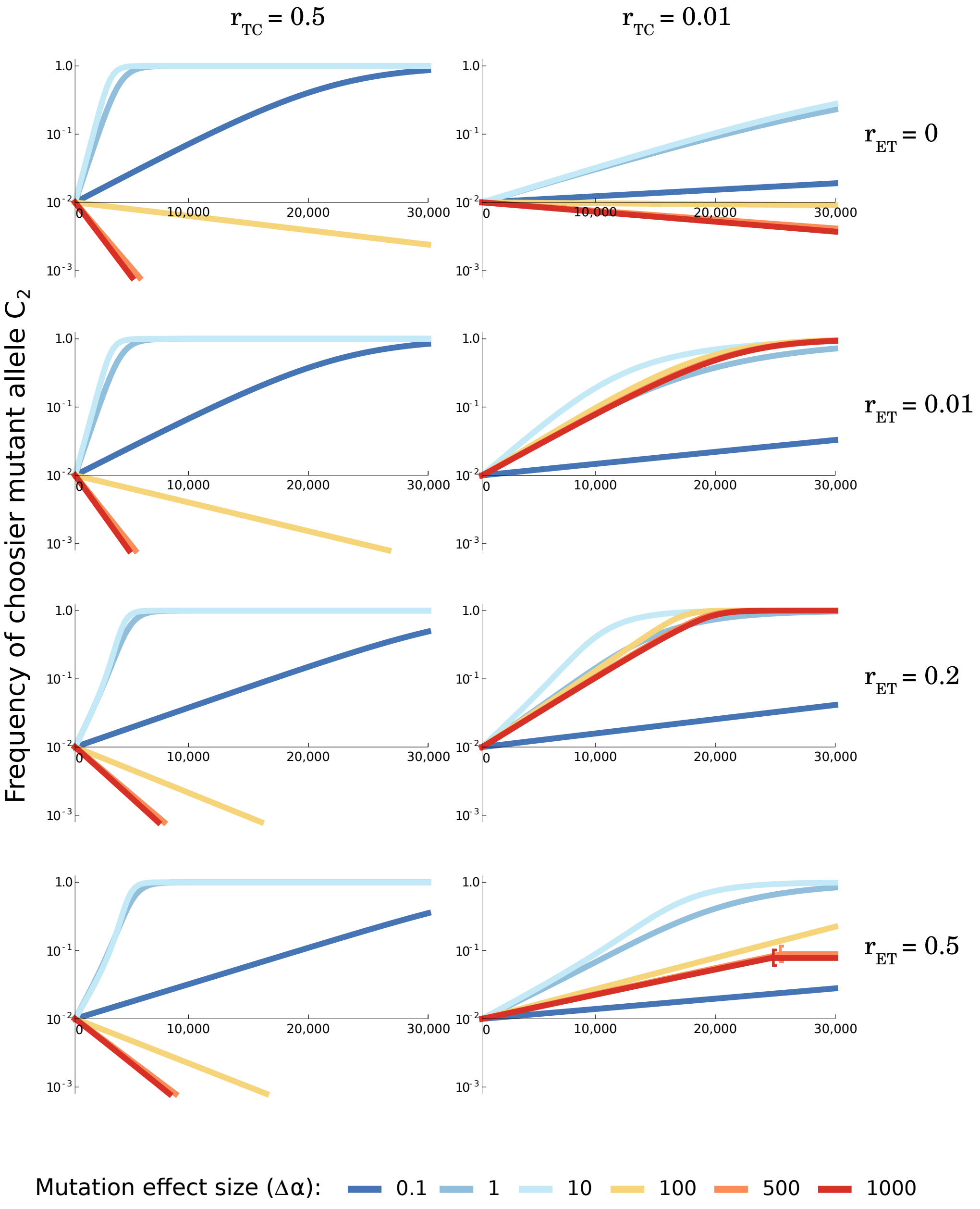


**Figure S14: Time series of the invasion of mutant alleles at the choosiness locus with a log-scale at the vertical axis.** Simulations are the same as in Figure 4.

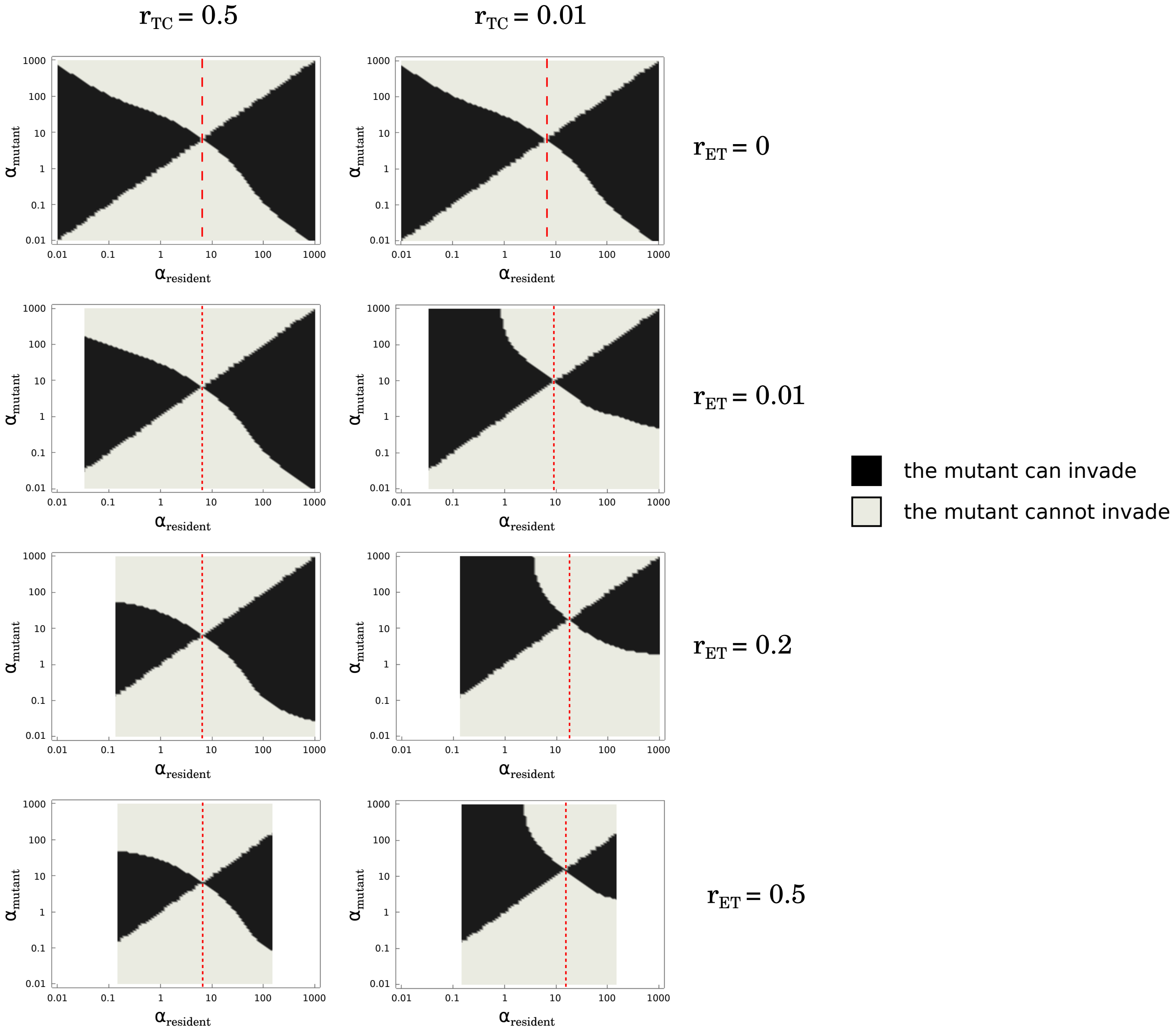


**Figure S15: Pairwise invasibility plots for different genetic architecture.** We represent whether a mutant allele (encoding choosiness $\alpha_{\text{mutant}}$) can invade in a resident population (with choosiness $\alpha_{\text{resident}}$) or not, when introduced at a frequency equal to 0.01. Our invasion criterion is based on whether or not the choosiness mutant allele increases in frequency over 200 generations after introduction. When polymorphism is lost at the T locus before introduction of the mutant allele, the mutant choosiness allele is neutral; this is represented by an uncolored zone (for extreme $\alpha_{\text{resident}}$ values when $r_{\text{ET}} > 0$). Note that we do not represent cases where the invasion of the mutant choosiness allele eventually leads to the loss of polymorphism (as shown in Fig. 4). These pairwise invasibility plots highlight that a nonmagic trait complex increases the ESS choosiness (dashed red line; as already shown in Fig. 2), and allows the evolution of very high choosiness through the spread of a single large-effect mutation which far overshoots the choosiness value at evolutionary equilibrium (for $r_{\text{TC}} = 0.01$; as shown in Fig. 4). Here, $m = 0.01$ and $s = 0.05$.

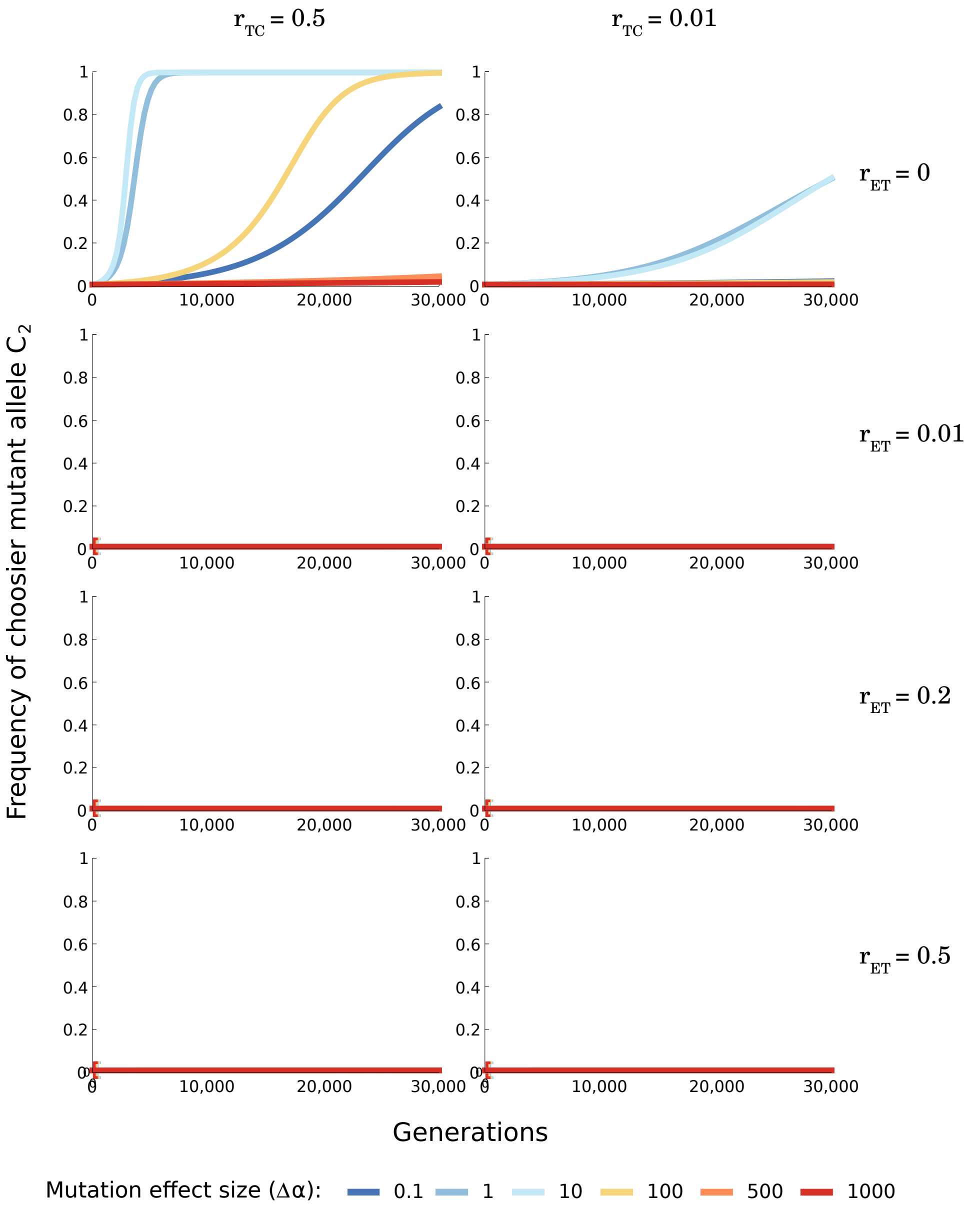


**Figure S16: Time series of the invasion of mutant alleles at the choosiness locus, when introduced in a population mating randomly at the onset of secondary contact.** We consider the same combination of parameters as in Fig. 4, but assume a different initial condition. We here initiate the populations in secondary contact (i.e., not at the selection-migration equilibrium) and we assume that mating is random initially ($\alpha_1 = 0$). Brackets show the time points where choosiness becomes a neutral trait because polymorphism at the T locus is lost (close to generation 0 in all panels for $r_{\text{ET}} > 0$). When $r_{\text{ET}}$ deviates from 0, polymorphism becomes lost at the onset of secondary contact, and choosiness has little chance to increase in frequency.

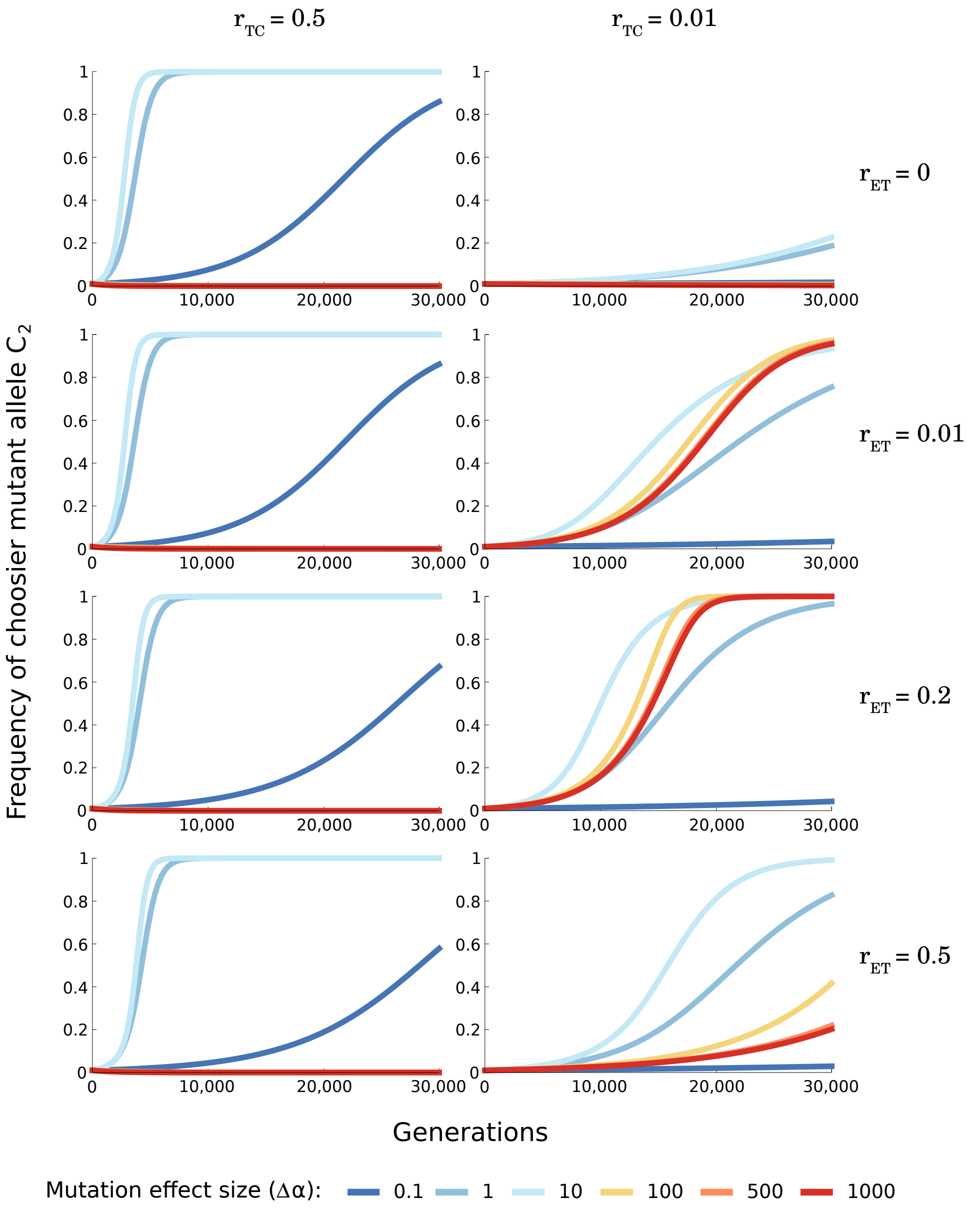


**Figure S17: Time series of the invasion of mutant alleles at the choosiness locus with asymmetric migration.** We consider the same combination of parameters as in Fig. 4, but assume asymmetric migration. Just like in Fig. 4, choosiness $\alpha_1$ is set to the choosiness value that maintains polymorphism at the T locus for all recombination rates tested ($\alpha_1 = 0.17$, estimated numerically; i.e., higher than in Fig. 4 because asymmetric migration inhibits the maintenance of polymorphism). Here, $m_1 = 0.01$, $m_2 = 0.011$, and $s = 0.05$. Polymorphism is lost for $r_{\text{ET}} = 0.5$ and $\Delta\alpha \geq 500$ after 30,000 generations (not shown here; but see Fig. S20 for time series lasting 100,000 generations). Here the initial condition differs from the one in Fig. 4, but see Fig S20 for a comparison with vs. without asymmetric migration starting from the same initial condition.

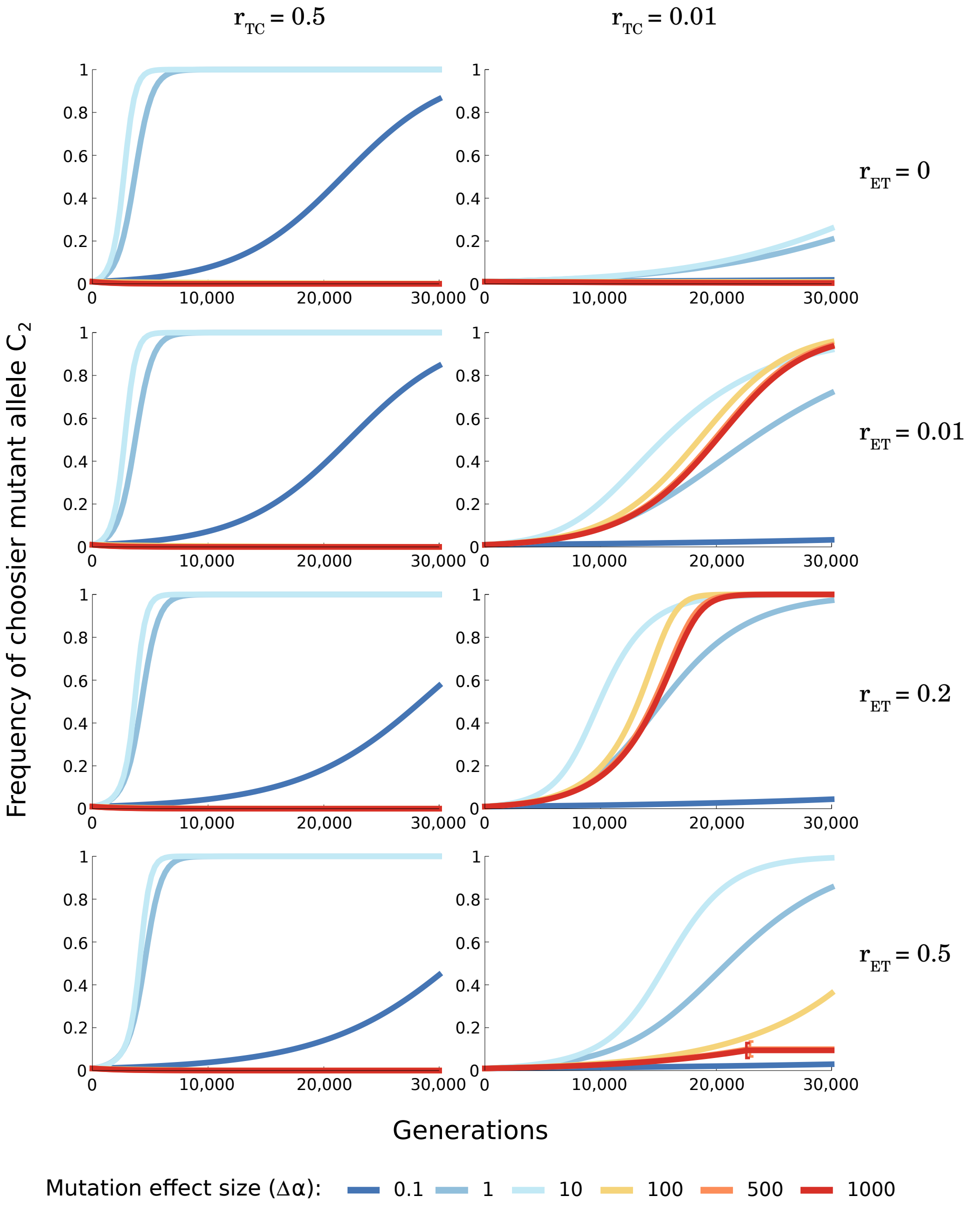


**Figure S18: Time series of the invasion of mutant alleles at the choosiness locus with asymmetric viability selection.** We consider the same combination of parameters as in Fig. 4, but assume asymmetric viability selection. Just like in Fig. 4, choosiness $\alpha_1$ is set to be the choosiness value that maintains polymorphism at the T locus for all recombination rates tested ($\alpha_1 = 0.14$, estimated numerically; i.e., higher than in Fig. 4 because asymmetric viability selection inhibits the maintenance of polymorphism). Brackets show the time points where choosiness becomes a neutral trait because polymorphism at the T locus is lost. Here, $m = 0.01$, $s_1 = 0.05$, and $s_2 = 0.055$. Just like with asymmetric migration, the loss of polymorphism, preventing the spread of a choosier allele, can occur more rapidly if viability selection is asymmetric, which is not surprising given that polymorphism is maintained under more constrained conditions with asymmetric viability selection than with symmetric viability selection (see Servedio and Bürger 2020, *Evolution* **74**(11):2438-2450). Here the initial condition differs from the one in Fig. 4, but see Fig S20 for a comparison with vs. without asymmetric viability selection starting from the same initial condition.

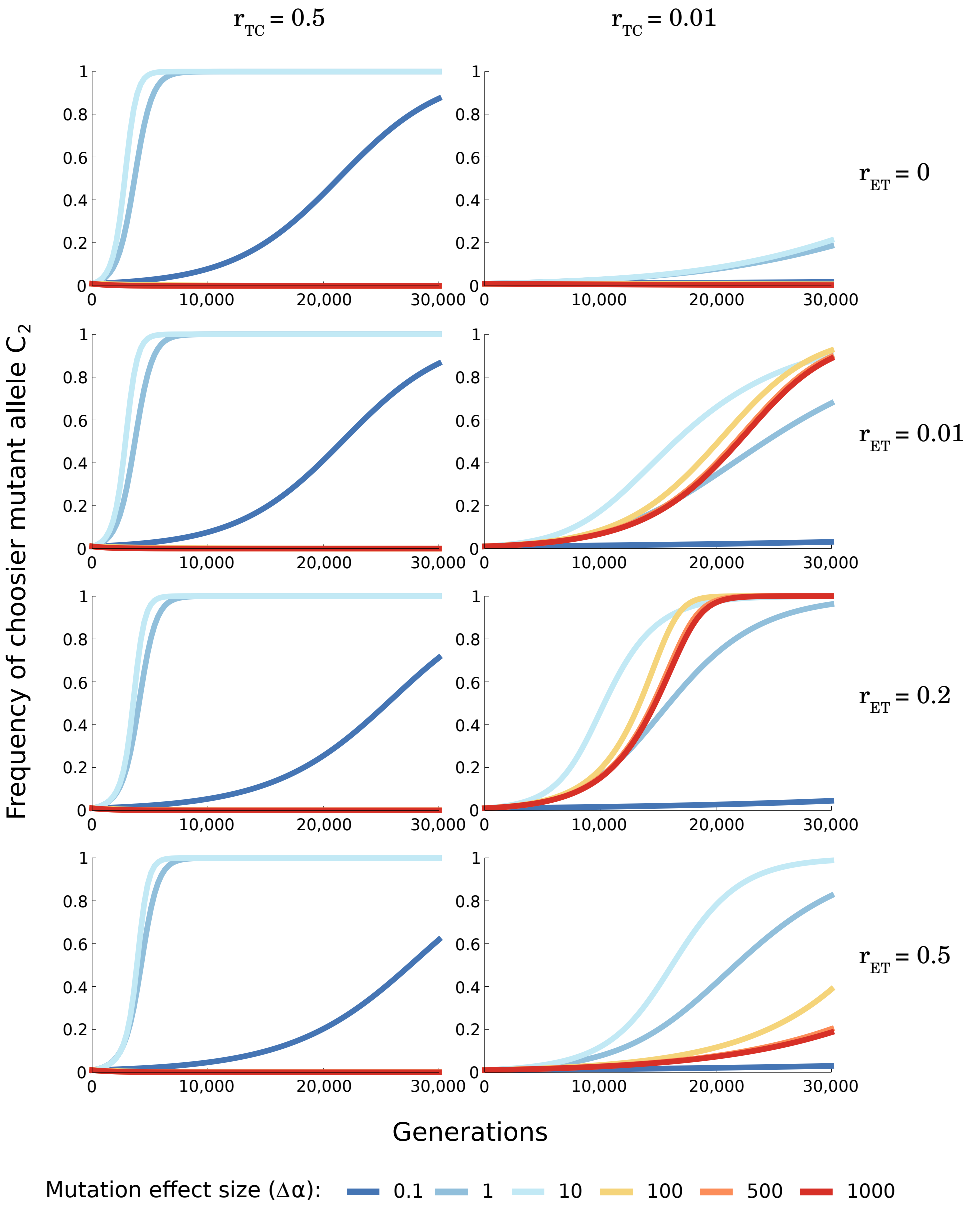


**Figure S19: Time series of the invasion of mutant alleles at the choosiness locus with asymmetric choosiness.** We consider the same combination of parameters as in Fig. 4, but assume asymmetric choosiness, with $T_2$ females having a 10% stronger choosiness than $T_1$ females do (with the allele being the same at the C locus). Just like in Fig. 4, choosiness $\alpha_1$ is set to be the choosiness value that maintains polymorphism at the T locus for all recombination rates tested ($\alpha_1 = 0.16$, estimated numerically; i.e., higher than in Fig. 4 because asymmetric choosiness inhibits the maintenance of polymorphism). Here, $m = 0.01$, and $s = 0.05$. Polymorphism is lost for $r_{\mathrm{ET}} = 0.5$ and $\Delta\alpha \geq 500$ after 30,000 generations (not shown here; but see Fig. S20 for time series lasting 100,000 generations). Here the initial condition (baseline $\alpha_1$ value) differs from the one in Fig. 4, but see Fig S20 for a comparison with vs. without asymmetric choosiness starting from the same initial condition.

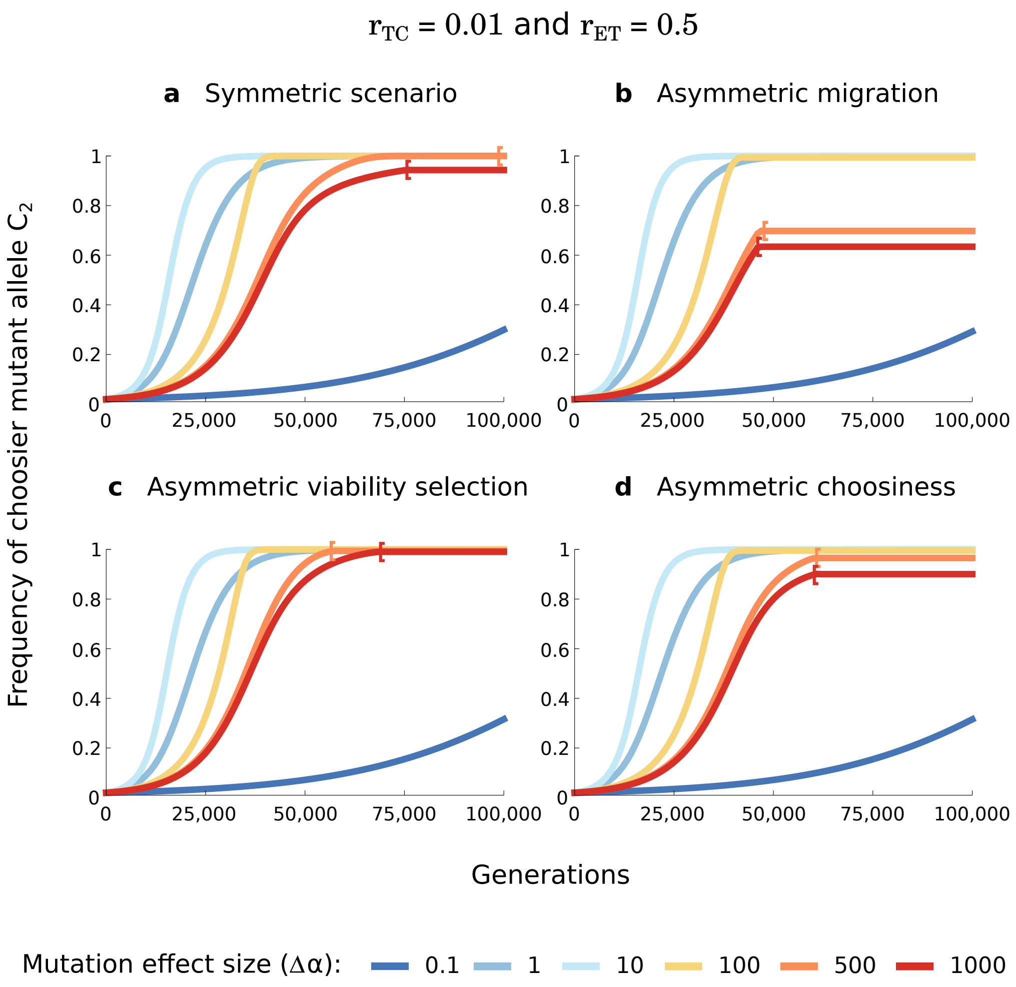


**Figure S20: Comparison of time series of the invasion of mutant alleles at the choosiness locus without (a) or with asymmetry (b-d).** We here show time series lasting 100,000 generations for $r_{\rm TC} = 0.01$ and $r_{\rm ET} = 0.5$ with symmetric conditions (**a**) or with some forms of asymmetry (**b**-**d**), in order to compare when polymorphism is lost. (**a**) Symmetric scenario with parameter values $m_1 = m_2 = 0.01$, $s_1 = s_2 = 0.05$, and with $T_1$ females having the same choosiness as $T_2$ females all else being equal. (**b**) Asymmetric migration with $m_1 = 0.01$ and $m_2 = 0.011$. (**c**) Asymmetric viability selection with $s_1 = 0.05$ and $s_2 = 0.055$. (**d**) Asymmetric choosiness with $T_2$ females having a 10% stronger choosiness than $T_1$ females do. In all cases, we implement $\alpha_1 = 0.17$ which allow the initial maintenance of polymorphism for all the conditions tested (we do not implement the same initial conditions in Figs. 4 and S17-S19 which gives the false impression that asymmetry prevents the loss of polymorphism). Brackets show the time points where choosiness becomes a neutral trait because polymorphism at the T locus is lost (polymorphism is not lost for $\Delta\alpha = 0.1$ even after 100,000 generations; not shown). Loss of polymorphism, preventing the spread of a choosier allele (in red and orange), can occur more rapidly under asymmetric conditions than under symmetric conditions, which is not surprising given that polymorphism is maintained under more constrained conditions with asymmetric conditions than with symmetric conditions (see Servedio and Bürger 2020, *Evolution* **74**(11):2438-2450 for more information on the conditions favoring the maintenance of polymorphism).

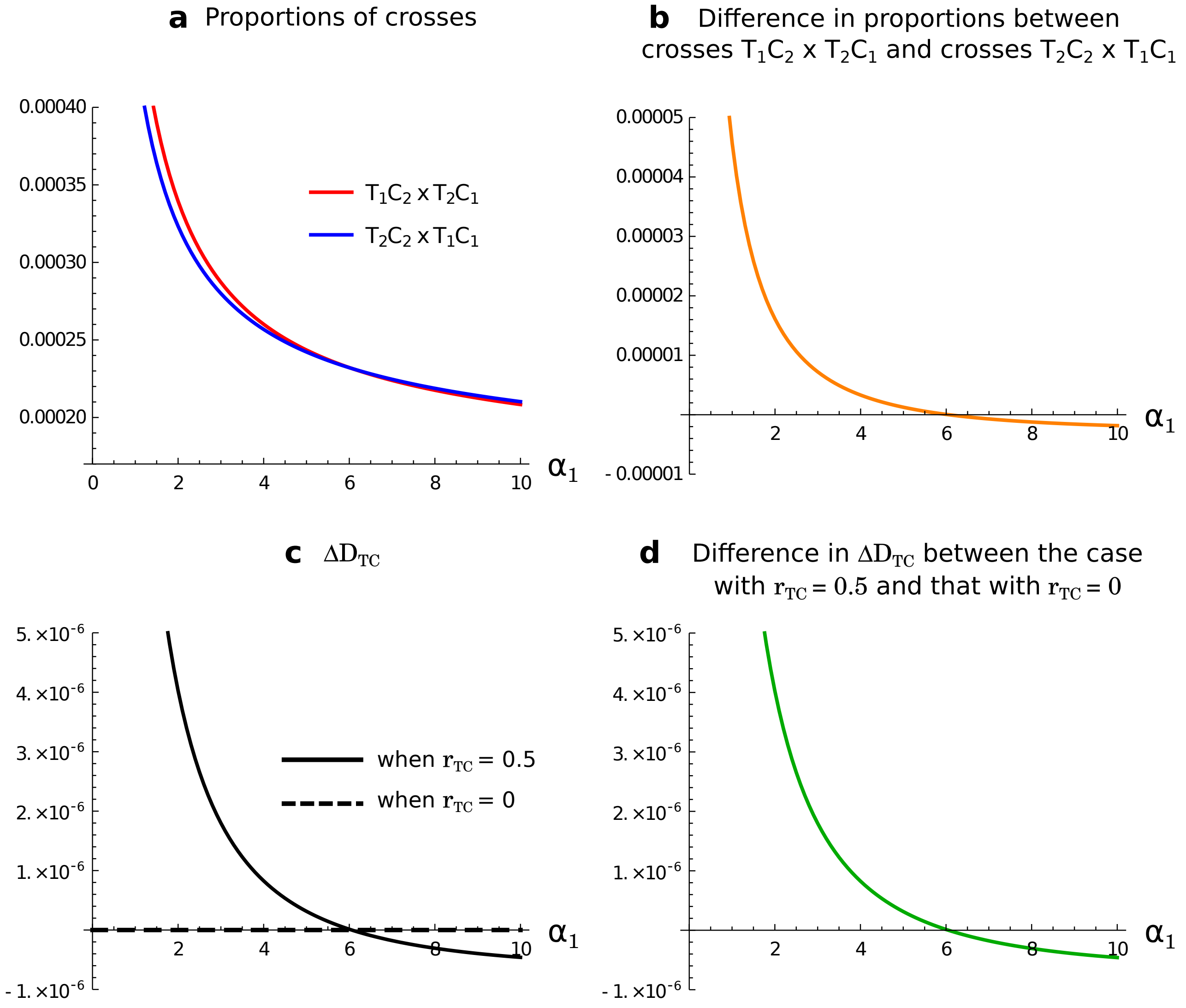


**Figure S21: Effect of the recombination rate between loci T and C on the build-up of linkage disequilibrium between loci T and C in the case of a magic trait.** We consider that allele $C_1$ codes for choosiness $\alpha_1$, and that allele $C_2$ codes for higher choosiness, $\alpha_2 = \alpha_1 + 1$. We start the population at the migration-selection equilibrium at the E and T loci given the ancestral choosiness, which is encoded by allele $C_1$. We introduce the allele $C_2$, coding for a higher choosiness, in linkage equilibrium with the other loci and in the same frequency (= 0.01) in both populations, and we allow migration and viability selection to occur. We then show, in population 2, (**a**) the proportions of crosses $T_1C_2 \times T_2C_1$ (red) and $T_2C_2 \times T_1C_1$ (blue) that respectively build and break the beneficial allelic association $T_2C_2$ via recombination between loci T and C, (**b**) the difference in proportions between these crosses, (**c**) the change in linkage disequilibrium, $\Delta D_{\text{TC}}$, during mating and the production of zygotes with or without recombination between loci T and C, and (**d**) the difference between these changes in linkage disequilibrium, $\Delta D_{\text{TC}}$. The lines in panel a, and the horizontal axis in panels **b-d**, are crossed at the value of $\alpha_1$ for which divergence is maximized. Here, $m = 0.01$, $s = 0.05$, and $r_{\text{ET}} = 0.0$.

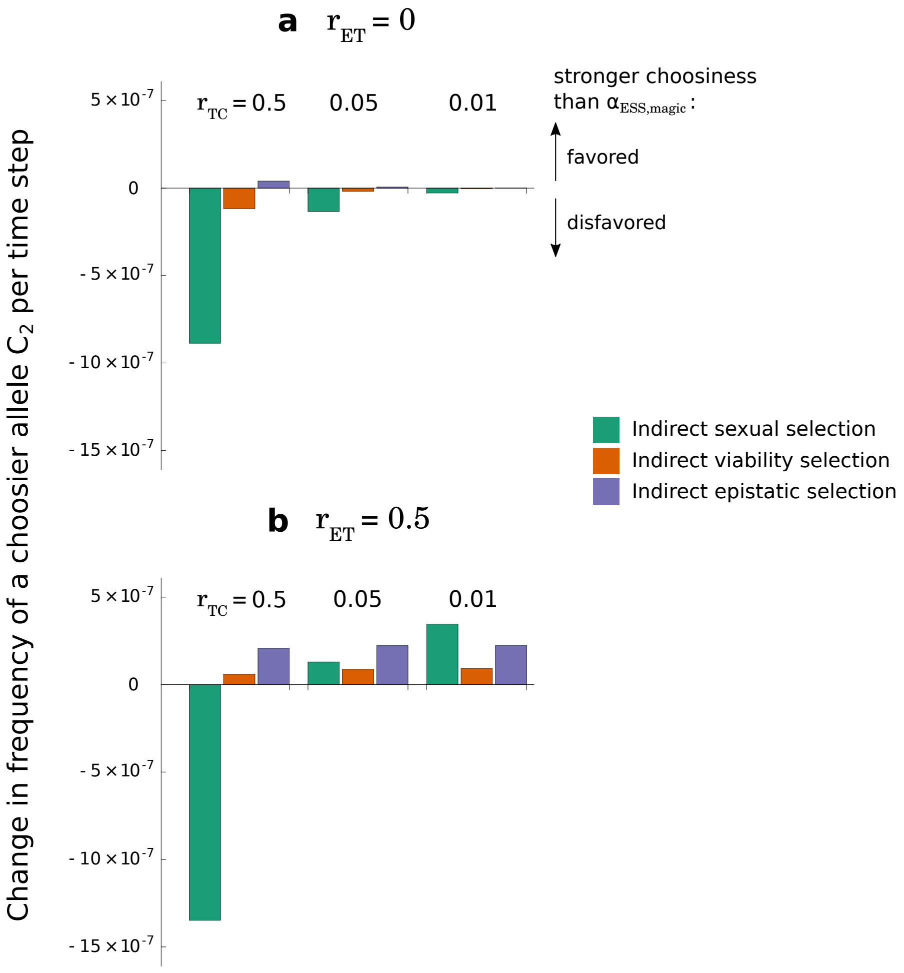


**Figure S22: Effect of the recombination rate between the choosiness locus and other loci, $\mathbf{r_{TC}}$, on selection acting on choosiness.** We consider the invasion of an allele coding for higher choosiness than the ESS choosiness for a magic trait such that $\alpha_1 = \alpha_{\text{ESS,magic}}$ and $\alpha_2 = \alpha_1 + 1$, and we represent first-order approximations of the effect of indirect sexual selection, indirect viability selection, and indirect epistatic selection on the change in frequency of the choosier allele (see caption of Fig. 3b-c for more details), for different recombination rates ($r_{\text{ET}}$ and $r_{\text{TC}}$). In panel **a**, we consider a magic trait ($r_{\text{ET}} = 0$ and maximum linkage disequilibrium between loci E and T), and in panel **b**, we consider a nonmagic trait ($r_{\text{ET}} > 0$). Here, $m = 0.01$, $s = 0.05$, and $\alpha_{\text{ESS,magic}} = 6.54$ (estimated numerically).
